# Anatomical and Functional Gradients Shape Dynamic Functional Connectivity in the Human Brain

**DOI:** 10.1101/2021.03.15.435361

**Authors:** Xiaolu Kong, Ru Kong, Csaba Orban, Wang Peng, Shaoshi Zhang, Kevin Anderson, Avram Holmes, John D. Murray, Gustavo Deco, Martijn van den Heuvel, B.T. Thomas Yeo

## Abstract

Large-scale biophysical circuit models can provide mechanistic insights into the fundamental micro-scale and macro-scale properties of brain organization that shape complex patterns of spontaneous brain activity. By allowing local synaptic properties to vary across brain regions, recent large-scale circuit models have demonstrated better fit to empirical observations, such as inter-regional synchrony averaged over several minutes, i.e. static functional connectivity (FC). However, most previous models do not capture how inter-regional synchrony patterns vary over timescales of seconds, i.e., time-varying FC dynamics. Here we developed a spatially-heterogeneous large-scale dynamical circuit model that allowed for variation in local circuit properties across the human cortex. We showed that parameterizing local circuit properties with both anatomical and functional gradients was necessary for generating realistic static and dynamical properties of resting-state fMRI activity. Furthermore, empirical and simulated FC dynamics demonstrated remarkably similar sharp transitions in FC patterns, suggesting the existence of multiple attractors. We found that time-varying regional fMRI amplitude tracked multi-stability in FC dynamics. Causal manipulation of the large-scale circuit model suggested that sensory-motor regions were a driver of FC dynamics. Finally, the spatial distribution of sensory-motor drivers matched the principal gradient of gene expression that encompassed certain interneuron classes, suggesting that heterogeneity in excitation-inhibition balance might shape multi-stability in FC dynamics.

## Introduction

Spontaneous fluctuations in large-scale brain activity exhibit complex spatiotemporal patterns across animal species (Hutchison et al., 2013; Gozzi and Schwarz, 2016; Ma et al., 2016; Betzel, 2020). Inter-regional synchrony of resting-state brain activity averaged over several minutes (i.e., time-averaged static functional connectivity) has informed our understanding of brain network organization (Damoiseaux et al., 2006; Smith et al., 2009; Gratton et al., 2018), individual differences in behavior (Finn et al., 2015; Kong et al., 2019) and mental disorders (Xia et al., 2018; Kebets et al., 2019). Recent studies have shown that additional important insights can be gained from studying moment-to-moment variation in inter-regional synchrony, i.e., time-varying dynamic functional connectivity (Allen et al., 2014; Zalesky et al., 2014; Vidaurre et al., 2017; Liegeois et al., 2019; Lurie et al., 2020). However, it is currently unclear how anatomical and functional heterogeneity in local circuit properties contribute to both time-averaged and time-varying properties of large-scale brain dynamics.

Large-scale spontaneous brain activity is thought to arise from the reverberation of intrinsic dynamics of local circuits interacting across long-range anatomical connections (Deco et al., 2011; Breakspear, 2017). Simulations of large-scale biophysically plausible models of coupled brain regions have provided mechanistic insights into spontaneous brain activity (Honey et al., 2007; Ghosh et al., 2008; Deco et al., 2014; Hansen et al., 2015). However, most previous large-scale circuit models assumed that local circuit properties (e.g., local synaptic strength, etc.) are identical across brain regions, which is not biologically plausible. Recent studies in both humans and macaques (Chaudhuri et al., 2015; Demirtas et al., 2019; Wang et al., 2019) have demonstrated that allowing local circuit properties to vary along the brain’s hierarchical axis yielded significantly more realistic static functional connectivity (FC). However, these heterogeneous models have not been shown to recapitulate time-varying FC dynamics.

In this study, we developed a spatially-heterogeneous mean field model (MFM) to realistically capture time-varying FC dynamics. Local circuit heterogeneity can be informed by *in-vivo* structural and functional neuroimaging measures. For example, T1-weighted/T2-weighted (T1w/T2w) MRI estimates of intracortical myelin and the principal resting-state FC gradient have been shown to index anatomical (Burt et al., 2018) and functional (Margulies et al., 2016) hierarchies respectively. Parameterization of local circuit properties with T1w/T2w maps led to more realistic static FC than a spatially-homogeneous mean field model (Demirtas et al., 2019). However, local circuit properties might be more strongly associated with the principal FC gradient than the T1w/T2w map (Wang et al., 2019). Thus, we hypothesized that parameterizing local circuit properties with both the T1w/T2w map and the principal FC gradient might lead to a more realistic computational model, which we will refer to as the parametric mean field model (pMFM). Using data from the Human Connectome Project (HCP), we demonstrated that pMFM achieved markedly more realistic static FC and FC dynamics in new out-of-sample participants, confirming the importance of functional and anatomical gradients to fully capture brain dynamics.

Both empirical and pMFM-simulated FC dynamics demonstrated remarkably similar sharp transitions in FC patterns, suggesting the existence of multiple FC states or attractors. Previous studies have suggested that multi-stability in nonlinear brain systems might arise from noise driven transitions between dynamic states or attractors (Freyer et al., 2012; Hansen et al., 2015; Deco et al., 2017). These noise-driven transitions might be reflected in the amplitude of regional brain activity. Therefore, we further investigated the relationship between the amplitude of regional fMRI signals and transitions in functional connectivity dynamics in both empirical and pMFM-simulated data. We also performed causal perturbations of the large scale circuit model to better understand the origins of FC multi-stability. Finally, the amplitude of regional fMRI signals have been linked with the gene expression markers of parvalbumin (PVALB) and somatostatin (SST) inhibitory interneurons (Anderson et al., 2020a), in line with rodent studies suggesting that differential interneuron abundance may underlie regional variability in local cortical function (Kim et al., 2017).

Thus, we also investigated the spatial relationship among FC dynamics, fMRI signal amplitude and gene expression patterns from the Allen Human Brain Atlas (AHBA). The contributions of this study are multi-fold. First, we showed that local circuit properties, parameterized by both anatomical and functional gradients, are important for generating realistic models of static FC and FC dynamics. Second, in both pMFM-simulations and empirical fMRI data, the amplitude of regional fMRI signals of sensory-motor regions tracked state transitions in FCD. Causal perturbations of the pMFM provide further evidence that sensory-motor regions might be drivers of FCD. Finally, the spatial distribution of sensory-motor drivers appeared to match the differential expression of PVALB and SST, as well as the first principal component of brain-specific genes. Overall, this suggests a potential link between FC dynamics and heterogeneity in excitation/inhibition balance across the cortex.

## Results

### Automatic optimization of the parametric mean field model (pMFM) yielded highly realistic functional connectivity dynamics

1052 participants from the HCP S1200 release were randomly divided into training (N = 351), validation (N = 350), and test (N = 351) sets. The Desikan-Killiany anatomical parcellation (Desikan et al., 2006) with 68 cortical regions of interest (ROIs) was used to generate group-averaged structural connectivity (SC) and static functional connectivity (FC) matrices from the training, validation and test sets separately. Analyses with a functional parcellation yielded similar conclusions (see “*Control analyses*”). For each rs-fMRI run, time-varying functional connectivity was computed using the sliding window approach (Allen et al., 2014; Liegeois et al., 2017). Briefly, for each rs-fMRI run, a 68 x 68 FC matrix was computed for each of 1118 sliding windows. Each window comprised 83 timepoints (or 59.76 seconds). The 68 x 68 FC matrices were then correlated across the windows, yielding a 1118 x 1118 functional connectivity dynamics (FCD) matrices for each run (Hansen et al., 2015; Liegeois et al., 2017).

The dynamic mean field model (MFM) was used to simulate neural dynamics of the 68 cortical ROIs (Deco et al., 2013). Based on the simulated neural activity at each ROI, the hemodynamic model (Stephan et al., 2007; Heinzle et al., 2016) was then used to simulate blood oxygen level–dependent (BOLD) fMRI. Details of the model can be found in the *Methods* section. Here we highlight the intuitions behind the MFM. In the MFM, the neural dynamics of each ROI are driven by four components: (1) recurrent (intra-regional) input, (2) inter-regional inputs, (3) external input (potentially from subcortical relays) and (4) neuronal noise. There are “free” parameters associated with each component. First, a larger recurrent connection strength *w* corresponds to stronger recurrent input current. Second, the inter-regional inputs depend on the neural activities of other cortical ROIs and the connectional strength between ROIs. The inter-regional connectional strength is parameterized by the SC matrices, scaled by a global scaling constant *G*. Third, *I* is the external input current. Fourth, the neuronal noise is assumed to be Gaussian with standard deviation *σ*.

In the current study, the recurrent connectional strength *w*, external input current *I*, and noise amplitude *σ* are each parameterized as a linear combination of the principal resting-state FC gradient (Margulies et al., 2016) and T1w/T2w myelin estimate (Glasser and Van Essen, 2011), resulting in 10 unknown linear coefficients. We refer to the resulting model as parametric MFM (pMFM). The 10 unknown linear coefficients were automatically estimated by minimizing disagreement between the empirical and simulated BOLD signal (Figure 1A).

**Figure 1.**
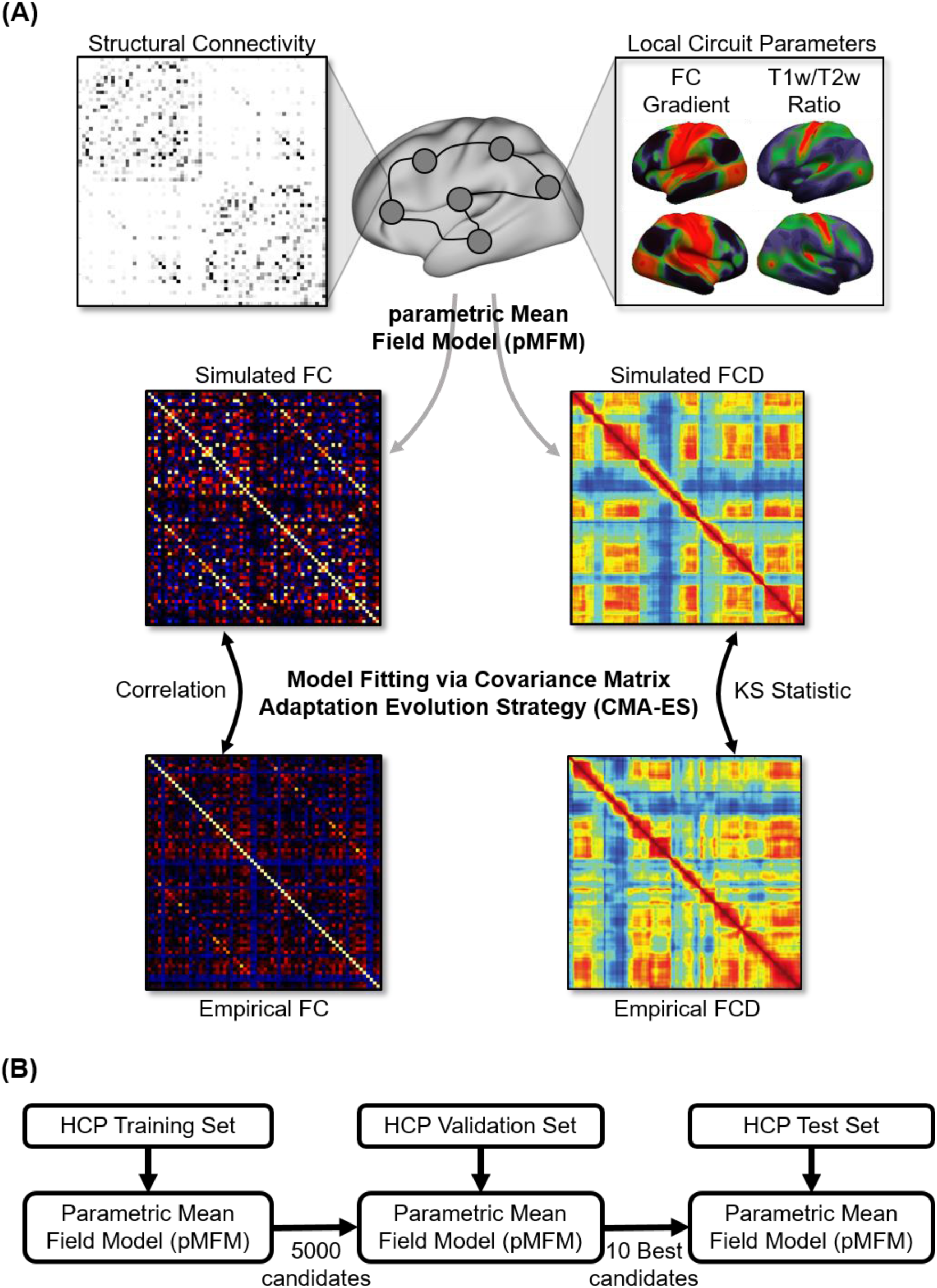
Schematic of parametric mean field model (pMFM) optimization. (A) The pMFM comprised ordinary differential equations (ODEs) at each cortical region coupled by a structural connectivity (SC) matrix. The circuit-level parameters were allowed to vary across cortical regions, parameterized by a linear combination of resting-state functional connectivity (FC) gradient and T1w/T2w spatial maps. The pMFM was used to generate simulated static FC and functional connectivity dynamics (FCD). The Covariance Matrix Adaptation Evolution Strategy (CMA-ES) was used to estimate the pMFM by minimizing a cost function of disagreement with empirically observed FC and FCD. (B) The CMA-ES algorithm was applied to the Human Connectome Project (HCP) training set (N = 351) to generate 5000 candidate parameter sets. The top 10 candidate parameter sets were then selected from the 5000 candidate sets based on the model fit in the validation set (N = 350). These top 10 candidate sets were then evaluated in the HCP test set (N = 351).

More specifically, the simulated fMRI was used to compute a 68 x 68 static FC matrix and a 1118 x 1118 FCD matrix. The agreement between the simulated and empirical static FC matrices was defined as the Pearson’s correlation (r) between the z-transformed upper triangular entries of the two matrices. Larger r indicated more similar static FC. The disagreement between the simulated and empirical FCD matrices was defined as the Kolmogorov–Smirnov (KS) distance between the upper triangular entries of the two matrices (Hansen et al., 2015). A smaller KS distance indicated more similar FCD. To optimize both static FC and FCD, an overall cost was defined as (1 - r) + KS and minimized in the training set. We considered three different minimization algorithms, each generating 5000 candidate sets of model parameters from the training set. Covariance matrix adaptation evolution strategy (CMA-ES; Hansen, 2006) performed the best in the validation set (Figure S1), so the 10 best CMA-ES parameter sets from the validation set were evaluated in the test set.

Figure 2A shows a representative empirical FCD from a participant in the test set. Figure 2B shows a simulated FCD generated by the pMFM using the best model parameters (from the validation set) using SC from the test set. Both empirical and simulated FCD exhibited red off-diagonal blocks representing recurring FC patterns. Across the 10 best candidate sets, KS distance between empirical and simulated FCD was 0.115 ± 0.031 (mean ± std). Correlation between empirical and simulated static FC was 0.66 ± 0.03. As a reference, the correlation between SC and static FC in the test set was 0.28.

**Figure 2.**
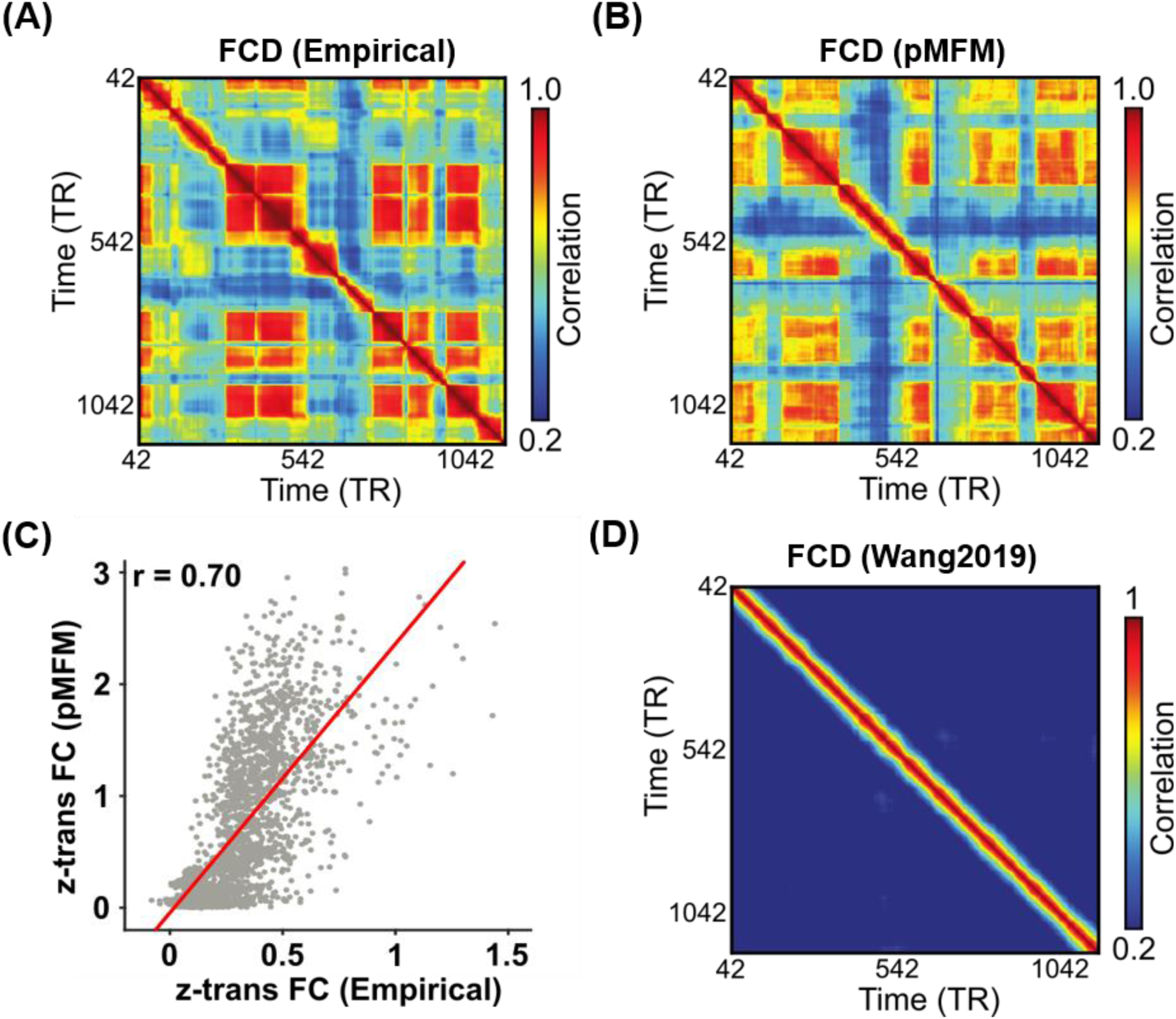
Parametric mean field model (pMFM) generates more realistic static functional connectivity (FC) and functional connectivity dynamics (FCD) than a previous spatially heterogeneous MFM (Wang et al., 2019). (A) Empirical FCD from a participant from the HCP test set. (B) Simulated FCD from the pMFM using the best model parameters from the validation set using structural connectivity (SC) from the test set. (C) Agreement (Pearson’s correlation) between empirically observed and pMFM-simulated static FC. (D) Simulated FCD generated by the previous spatially heterogeneous MFM (Wang et al., 2019).

Figure 2C shows the simulated FCD using the MFM parameters from our previous study (Wang et al., 2019). The almost constant values in off-diagonal elements suggests a lack of realistic FC dynamics. KS distance between empirical and simulated FCD was 0.88. Correlation between static empirical and simulated static FC was 0.48. Thus, the pMFM was able to generate much more realistic static FC and FCD than the MFM (Wang et al., 2019).

### Anatomical & functional gradients are critical to generating functional connectivity dynamics

In the previous section, we demonstrated that pMFM was able to generate realistic static FC and FCD. To explore what aspects of pMFM are important for generating realistic static FC and FCD, we performed a number of control analyses. First, we investigated the importance of utilizing both anatomical and functional gradients in generating realistic static FC and FCD. Most large-scale circuit model studies assume spatially homogeneous parameters. When recurrent connectional strength *w*, external input current *I*, and noise amplitude *σ* were optimized by CMA-ES, but constrained to be spatially homogeneous (Figure 3), then there was substantially weaker agreement with empirical static FC (r = 0.56 ± 0.05) and FCD (KS = 0.50 ± 0.30). Similarly, spatial heterogeneity for all three parameters (*w*, *I* and *σ*) were necessary to generate the most realistic static FC and FCD in the test set (Figures S2A to S2C).

**Figure 3.**
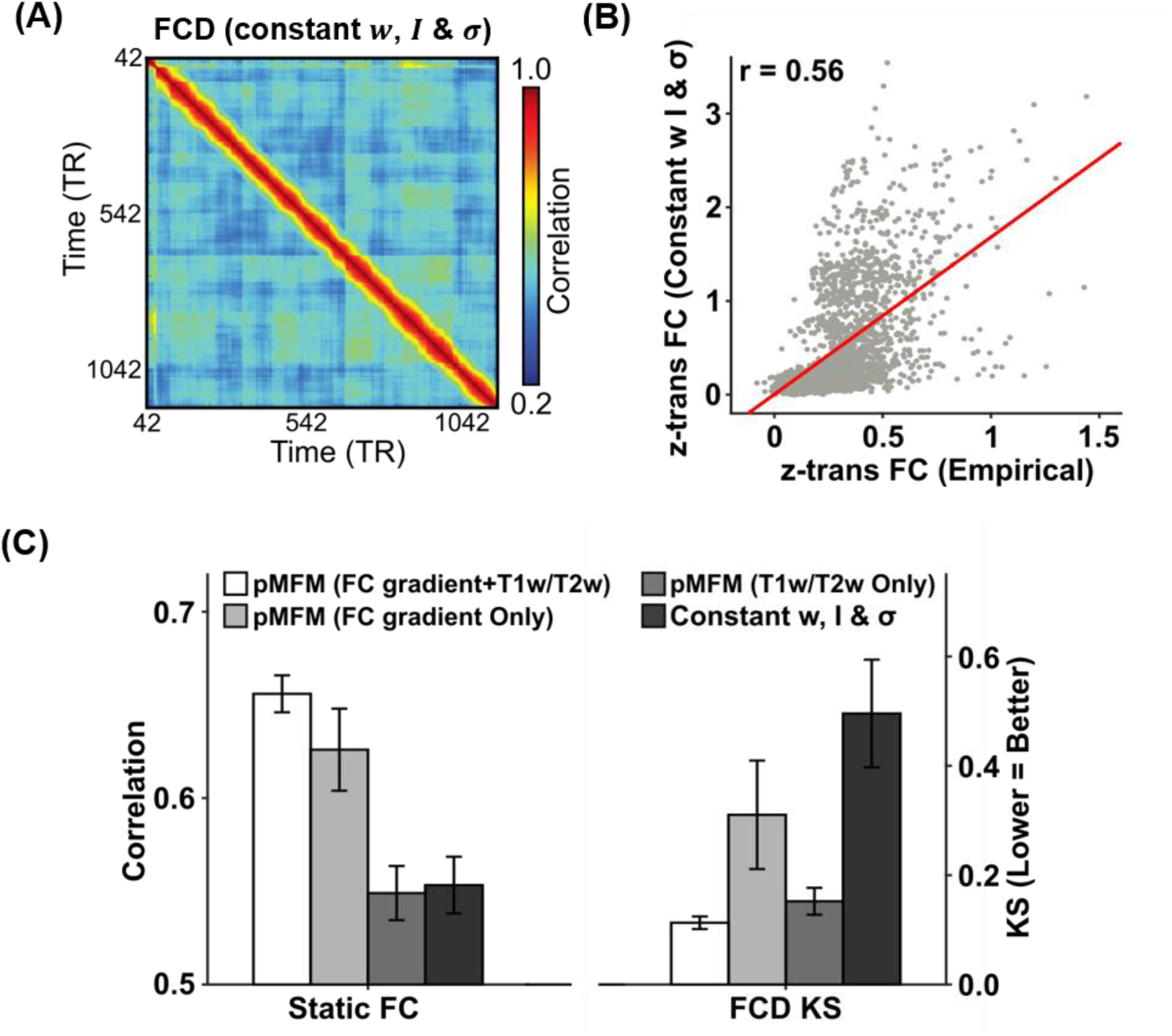
Importance of multiple spatial gradients for generating realistic static functional connectivity (FC) and functional connectivity dynamics (FCD). (A) Simulated FCD from a mean field model (MFM) optimized using the same algorithm as pMFM, but with model parameters constrained to be the same across cortical regions. (B) Agreement between empirically observed and simulated static FC from MFM optimized using the same algorithm as pMFM, but with model parameters constrained to be the same across cortical regions. (C) Agreement (Pearson’s correlation) between simulated and empirically observed static FC, as well as disagreement (KS distance) between simulated and empirically observed FCD across different conditions. The pMFM utilizing both anatomical and functional gradients (FC gradient and T1w/T2w spatial maps) performed the best, suggesting that T1w/T2w and FC gradient provided complementary contributions.

Second, if recurrent connectional strength *w*, external input current *I*, and noise amplitude *σ* were parameterized with only T1w/T2w (i.e., Demirtas et al., 2019) or only FC gradient, then the resulting static FC and FCD were less realistic in the test set (Figure 3C). Furthermore, if recurrent connectional strength *w*, external input current *I*, and noise amplitude *σ* were allowed to be spatially heterogeneous across brain regions, but not constrained by T1w/T2w or FC gradient (i.e., non-parametric), then simulations could achieve realistic static FC, but not FCD (Figure S2D). One reason could be the large number of “free” parameters leading to overfitting in the training set.

Finally, instead of fitting to both static FC and FCD in the training set, we also tried fitting only to static FC. Not surprisingly, the resulting model yielded unrealistic functional connectivity dynamics (Figure S3; KS = 0.88 ± 0.004). On the other hand, correlation between static empirical and simulated static FC was 0.74 ± 0.01, which was only slightly better than when optimizing both static FC and FCD (Figure 2C). This suggests that the goals of generating realistic static FC and FCD were not necessarily contradictory.

Overall, these results suggest the importance of parameterizing recurrent connectional strength *w*, external input current *I*, and noise amplitude *σ* with spatial gradients that smoothly varied from sensory-motor to association cortex. Furthermore, T1w/T2w and FC gradient are complementary in the sense that combining the two spatial maps led to more realistic static FC and FCD (Figure 3).

### Opposite gradient directions in recurrent connection strength, noise amplitude and external input

Figures 4B to 4D illustrate the spatial distribution of recurrent connection strength *w*, external input current *I*, and noise amplitude *σ* based on the best parameter estimate from the validation set. The black lines indicate seven resting-state network boundaries (Figure 3A; Yeo et al., 2011). While the resting-state network boundaries do not exactly align with the anatomically defined parcels, there was a striking correspondence between the resting-state networks and estimated pMFM parameters. Given the parameterization of pMFM by a linear combination of FC gradient (Margulies et al., 2016) and T1w/T2w spatial maps (Demirtas et al., 2019), it was not surprising that the parameter estimates exhibited a hierarchical gradient of values monotonically changing from sensory-motor to association networks (right column of Figures 4B to 4D).

**Figure 4.**
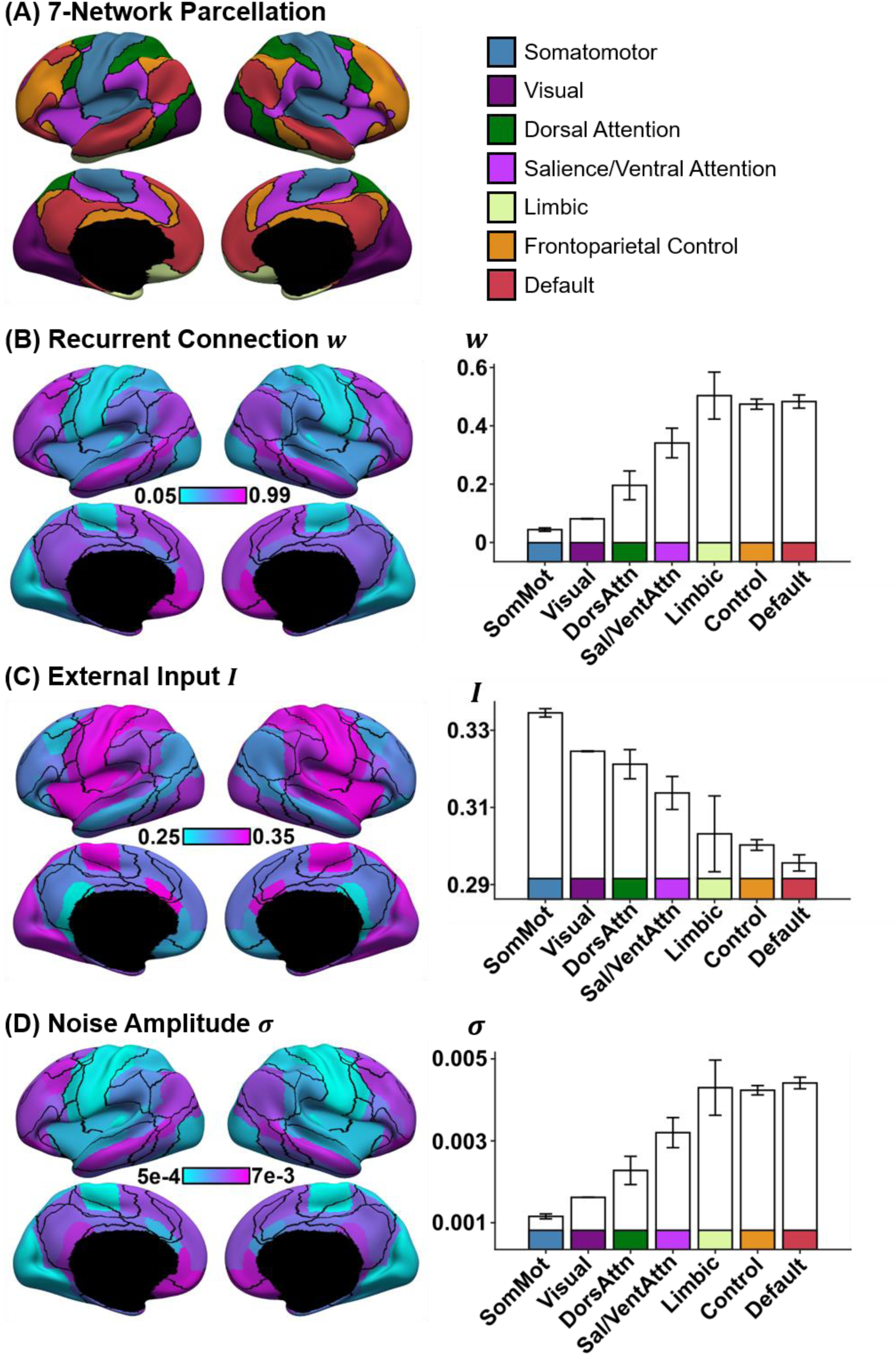
Spatial distribution of recurrent connection strength *w*, external input current *I*, and noise amplitude *σ*, and their relationships with resting-state networks. (A) Seven resting-state networks (Yeo et al., 2011). (B) Strength of recurrent connection *w* in 68 Desikan-Killiany cortical ROIs (left) and seven resting-state networks (right). (C) Strength of external input *I* in 68 Desikan-Killiany cortical ROIs (left) and seven resting-state networks (right). Strength of noise amplitude *σ* in 68 Desikan-Killiany cortical ROIs (left) and seven resting-state networks (right). The bars represent the mean values across regions within each network. The error bars show the standard error across regions within each network. Recurrent connection strength and noise amplitude increased from sensory-motor to association (limbic, control and default) networks. On the other hand, external input current was the highest in sensory-motor networks and decreased towards the default network.

However, the gradient directions were different across the three parameters. In particular, both recurrent connection strength and noise amplitude appeared to increase from sensory-motor to association (limbic, control and default) networks. On the other hand, external input current was the highest in sensory-motor networks and decreased towards the default network. The directionalities of noise amplitude and external input current were consistent across all the top ten parameter estimates from the validation set. In the case of recurrent connection strength, one of the ten parameter sets exhibited the opposite direction (i.e., decrease from sensory-motor regions to association networks; Figure S4), suggesting potential degeneracy in the case of recurrent connection strength.

### Time-varying amplitude of regional fMRI time courses tracks time-varying functional connectivity

Given that the pMFM was able to generate realistic FCD, we now seek to use the pMFM to provide further insights into mechanisms underlying FCD. Previous studies have suggested that FCD might arise from switching between multi-stable states (Hansen et al., 2015; Deco et al., 2017). Indeed, a magnified portion of the FCD matrix from a HCP test participant (Figure 5A) suggests the presence of at least two distinct states. In one state (white asterisk in Figure 5A), the sliding window FC pattern appeared to be coherent over a period of time. In a second state (black asterisk in Figure 5A), the sliding window FC patterns were incoherent over a period of time, so the high correlations within the block were restricted to the diagonals, and likely driven by autocorrelation in the fMRI signals and overlapping sliding windows. We hypothesized that fMRI signals might be dominated by large coherent amplitude fluctuations during the coherent state and dominated by noise during the incoherent state (right panel in Figure 5A; see Cocchi et al., 2017 for a review of multi-stability). If our hypothesis were true, we would expect large regional fMRI signal amplitude during the coherent state and small regional fMRI signal amplitude during the incoherent state.

**Figure 5.**
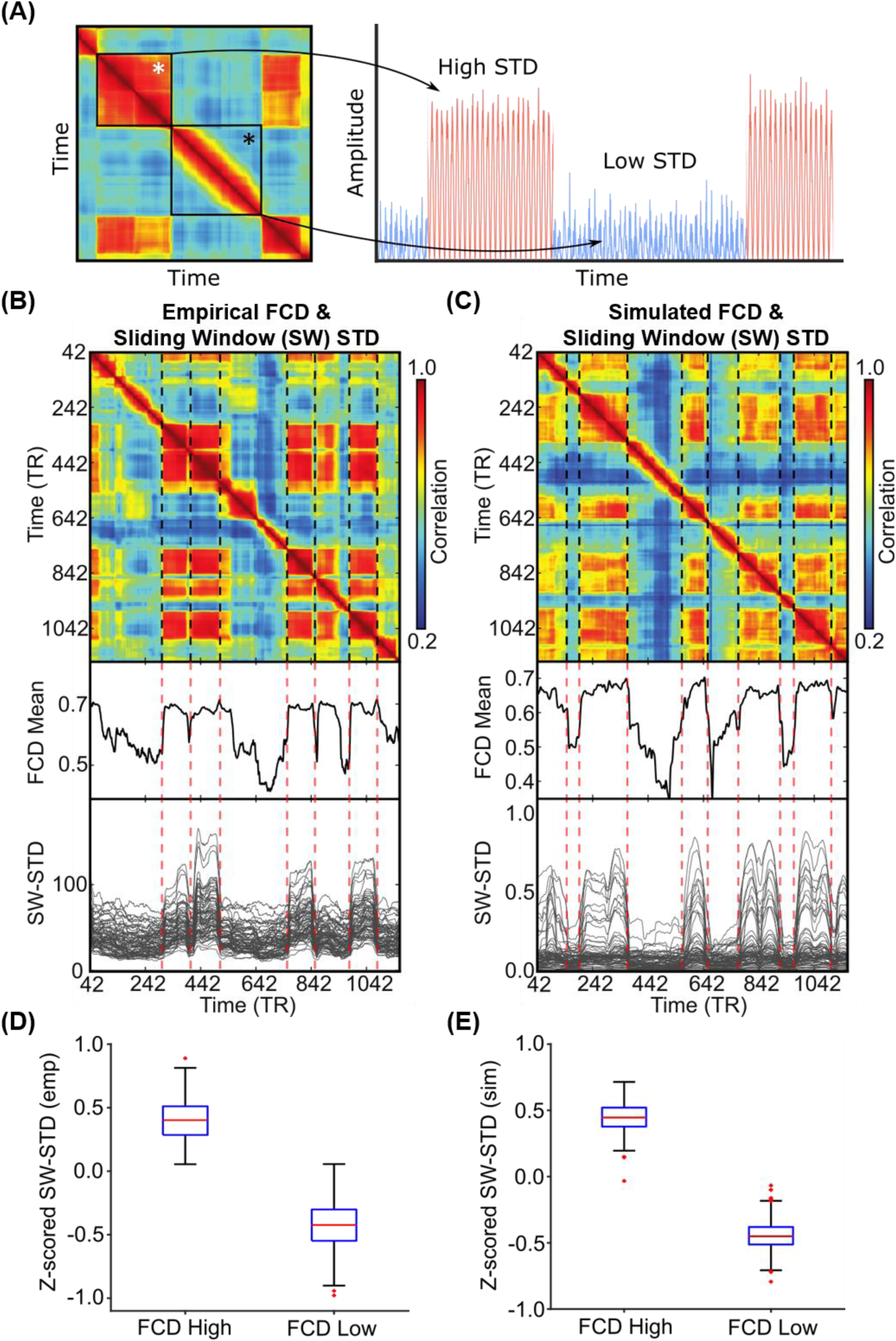
Correspondence between functional connectivity dynamics (FCD) and time-varying amplitude of regional fMRI time courses. (A) Inspection of FCD from a HCP test participant suggests at least two states. The first state (white asterisk) exhibits coherent FC patterns over a period of time. The second state (black asterisk) exhibits incoherent FC patterns over a period of time. The right panel illustrates our hypothesis that the coherent state might be characterized by large coherent amplitude in regional fMRI signals, i.e., high standard deviation (STD), while the incoherent state might be characterized by noise in regional fMRI signals, i.e., low standard deviation (STD). (B) Top panel shows empirical FCD matrix of a HCP test participant. The middle panel shows the FCD mean time course obtained by averaging the rows of the FCD matrix from the top panel. The bottom panel shows the standard deviation of each regional fMRI time course within each sliding window (SW- STD). The color of the lines corresponds to the correlation between the first derivative of the FCD mean time course and the first derivative of the SW-STD time courses. Sharp transitions in SW-STD corresponded to sharp FCD transitions (red dashed lines). (C) Same as panel B, but simulated from pMFM using the best model parameters from the validation set and structural connectivity from the test set. (D) SW-STD during coherent (high FCD mean) and incoherent (low FCD mean) states. Boxplots illustrate variation across HCP test participants. Coherent states were characterized by large amplitude (STD) in fMRI signals (p = 2.4e-168). (E) Same as panel D, but simulated from pMFM.

To test our hypothesis, the standard deviation of average fMRI signal of each cortical ROI within each sliding window was computed. Figure 5B (top panel) shows the FCD matrix of a single participant from the HCP test set. Figure 5C (top panel) shows the simulated FCD matrix from the pMFM using the best model parameters from the validation set and structural connectivity (SC) from the test set. The middle panels of Figures 5B and 5C show the FCD mean time course obtaining by averaging the rows of the FCD matrices from the top panels. Sharp transitions in the FCD mean time course reflected sharp transitions in the FCD matrix. The bottom panel shows the sliding window standard deviation (SW-STD) of empirical and simulated fMRI signals. There was striking correspondence between sharp transitions in the FCD mean time course and SW-STD time courses in both empirical and simulated data (red dashed lines in Figures 5B and 5C).

Consistent with our hypothesis, there was large signal amplitude during the coherent state and low signal amplitude during the incoherent state (Figure 5B). To quantify this phenomenon, for each run of each participant in the HCP test set, the top 10% of each FCD mean time course was designated as the coherent state (high FCD mean) and the bottom 10% of each FCD mean time course was designated as the incoherent state (low FCD mean). The SW-STD was then averaged across all cortical regions and across all runs of each participant. As shown in Figure 5D, the SW-STD was significantly higher during the coherent state than the incoherent state (p = 2.4e-168). Similar results were obtained for the pMFM simulations (Figure 5E).

### Sensory-motor regions drive switching behavior in functional connectivity dynamics

In the previous section, we found striking correspondence between the FCD mean time course and the regional SW-STD time courses (Figures 5B & 5C). We note that the FCD mean time course reflected cortex-wide fluctuations in FC patterns, while SW-STD time courses were region-specific. Therefore, to investigate regional heterogeneity of FCD-STD correspondence (Figure 5) across the cortex, correlation between the first derivative of the FCD mean time course and the first derivative of the SW-STD time course was computed for each cortical region. In the case of empirical observations, the FCD-STD correlations were averaged across all runs of all participants in the test set yielding a final FCD-STD correlational spatial map (Figure 6A). In the case of pMFM simulations, the correlations were averaged across 1000 random simulations using the best model parameters from the validation set using structural connectivity (SC) from the test set, yielding a final FCD-STD correlational spatial map (Figure 6B).

**Figure 6.**
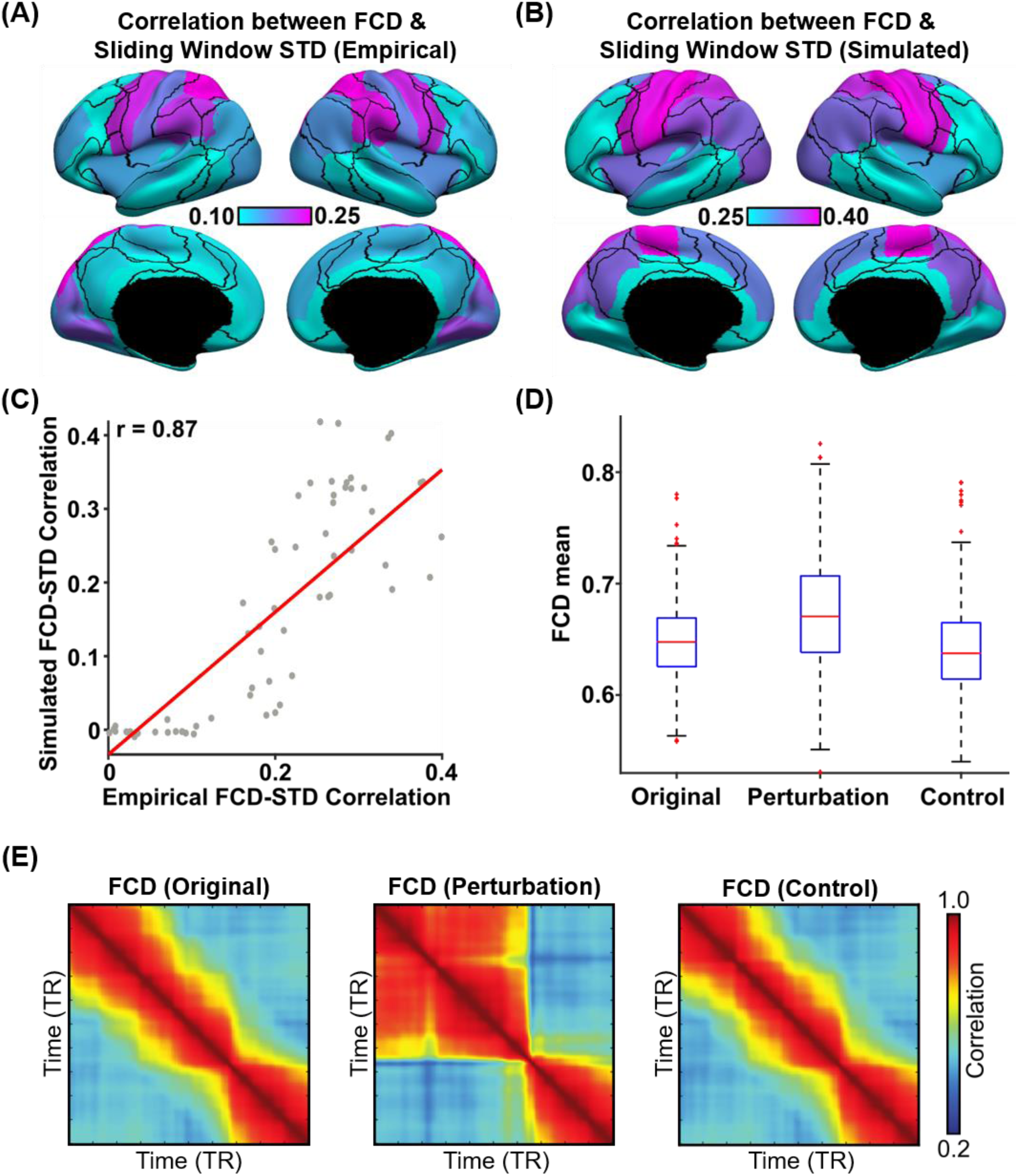
Sensory-motor regions drive sharp transitions in functional connectivity dynamics (FCD). (A) FCD-STD correlations obtained by correlating the first derivative of the FCD mean time course and the first derivative of the SW-STD time course of each cortical region. These correlations were performed for each HCP test participant and averaged across all runs and participants. (B) Same as panel A but simulated from pMFM using the best model parameters from the validation set and structural connectivity from the test set. The correlations were averaged across 1000 random simulations. (C) Correlation between empirical and simulated FCD-STD correlation spatial maps from panels B and C, showing strong correspondence between empirical and simulated results. (D) Casual perturbation of top 5 FCD-STD correlated regions (panel B) during the incoherent state (low FCD mean) led to transition into the coherent state (high FCD mean). As a control analysis, perturbation of the bottom 5 FCD-STD correlated regions (panel B) during the incoherent state (low FCD mean) did not lead to a state change (FCD mean remains low). (E) Example FCD from the perturbation experiments. (Left) original incoherent state. (Middle) perturbation of top 5 FCD-STD correlated regions (sensory-motor drivers). (Right) perturbation of bottom 5 FCD- STD correlated regions.

Statistical significance was established using a permutation test (see Methods). Almost all cortical regions were significant after correcting for multiple comparisons (FDR q < 0.05; Figure S5). Across both pMFM simulations and empirically observed data, FCD-STD correlations were the highest in sensory-motor regions and lowest in association cortex. There was strong spatial correspondence between simulated and empirical results (r = 0.87; Figure 6C). We note that the pMFM was optimized to yield realistic FCD with no regard for spatial correspondence, so the high level of spatial correspondence suggests that the pMFM was able to generalize to new unseen properties of FCD.

To explore the causal relationship between sensory-motor regions and FCD, we tested whether perturbation of sensory-motor regions could “kick” the system from an incoherent FCD state to a coherent FCD state. Among 1000 random simulations of pMFM, time segments in the incoherent state (low FCD mean) lasting for at least 200 contiguous fMRI timepoints were selected. The neural signals of the top five FCD-STD regions (sensory-motor drivers; Figure 6B) were then perturbed to increase their amplitude. The perturbation led to the successful transition of the FCD into a more coherent state with higher FCD mean (p = 6e-14; Figure 7D). Perturbation of the bottom five FCD-STD regions (Figure 6B) did not lead to an increase in FCD mean. Figure 7E illustrates example results of the perturbation experiment. Similar results were obtained if we perturbed top 10 and bottom 10 regions. Overall, this suggests that sensory-motor regions were a driver of switching behavior in FCD.

**Figure 7.**
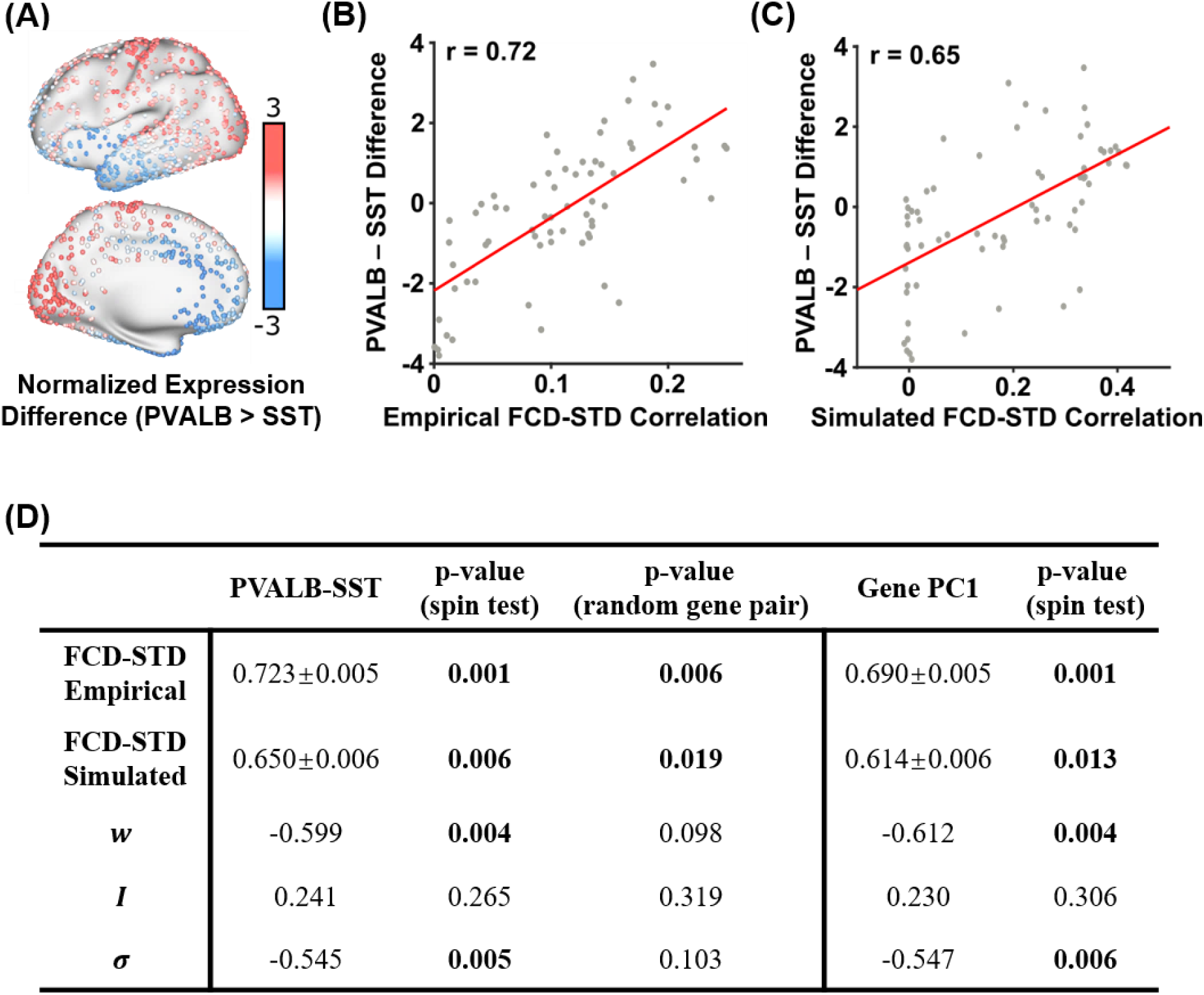
Correlations between the spatial distribution of sensory-motor drivers (FCD-STD correlational spatial maps) and gene expression spatial maps. (A) Difference in normalized expressions of parvalbumin and somatostatin (PVALB-SST) from the Allen Human Brain Atlas (AHBA). Panel is a re-rendering of (Anderson et al. 2020a). (B) Correlation between empirical FCD-STD correlational map (Figure 6B) and PVALB-SST gene expression map. (C) Correlation between simulated FCD-STD correlational map (Figure 6C) and PVALB/SST gene expression map. (D) Table of correlations between FCD-STD correlational spatial maps and two gene expression maps: PVALB-SST and first principal component of gene expression (Burt et al., 2018; Anderson et al., 2020b). The “spin test” tested the significance of the correlations while controlling for spatial autocorrelation. The “random gene pair” tested for the specificity of PVALB-SST by randomly sampling pairs of brain-specific genes. P values that survived the false discovery rate (q < 0.05) are bolded. Standard deviations reported in the table were obtained by bootstrapping.

### Parvalbumin-somatostatin and first genetic principal component correlate with sensory-motor drivers of time-varying functional connectivity dynamics

Results from the previous sections suggest that time-varying amplitude of sensory-motor regions tracks switching behavior in time-varying functional connectivity. A recent study (Anderson et al., 2020a) demonstrated that difference in the spatial distribution of molecular markers of parvalbumin and somatostatin interneurons (PVALB-SST) is linked with the amplitude of regional fMRI signals (Figure 7A). This intriguing finding is in line with data in rodents documenting the importance of these interneuron classes in local cortical circuit function (Kim et al., 2017). Inspection of the cortical distribution of PVALB-SST transcripts from the Allen Human Brain Atlas (AHBA) dataset (Figure 7A) suggests strong similarity with the FCD-STD correlational spatial maps (Figure 6).

PVALB -SST (Figure 7A) was averaged within each cortical ROI and correlated with the FCD-STD correlational spatial maps (Figure 6). The correlations were 0.72 and 0.65 for the empirical (Figure 7B) and simulated (Figure 7C) data respectively. As shown in Figure 7D, both correlations were significant based on spin-tests preserving spatial autocorrelation (Gordon et al., 2016; Alexander-Bloch et al., 2018). To test for specificity of PVALB-SST, a null distribution was also generated based on random pairs of brain-specific genes. Both correlations were again significant (Figure 7D). Overall, this suggests that the spatial distribution of sensory-motor drivers was associated with the differential expression of PVALB and SST

Given that previous studies have suggested the existence of multiple similar gene expression gradients, the first principal component of AHBA brain-specific gene expression data (Burt et al., 2018; Anderson et al., 2020b) was correlated with the FCD-STD correlational spatial maps (Figure 6). The first gene expression principal component was also correlated with both empirical and simulated FCD-STD spatial maps, although the correlations were slightly weaker than the correlations with PVALB-SST gene expression map (Figure 7D).

The recurrent connection strength *w* and noise amplitude *σ* were also correlated with the PVALB-SST gene expression map under the spin-test, but not the random-gene-pair tests. This suggests a lack of specificity to PVALB-SST (Figure 7D). The external input *I* was not correlated with any gene expression pattern.

### Control analyses

To ensure robustness of results, we performed several control analyses. First, we note that the simulation of pMFM utilized 10ms time step. To ensure that this time step was sufficiently small, the best model parameters from the validation set was applied to the test set using 1ms time step. KS distance between empirical and simulated FCD in the test set was 0.113 ± 0.047. Correlation between empirical and simulated static FC was 0.344 ± 0.033.

Second, the previous analyses utilized sliding window comprising 83 timepoints for computing FCD. To ensure the model parameters generalized to different window lengths, empirical and simulated FCD was computed in the test set using window lengths of 43 and 125. KS distance between empirical and simulated FCD in the test set was 0.148 ± 0.068 and 0.67 ± 0.040 for window lengths 43 and 125 respectively.

Third, we investigated whether the FCD-STD correlation maps (Figure 6) might be influenced by global signal fluctuation. We repeated the analysis by restricting to 50 test participants with the lowest global signal fluctuation. The resulting FCD-STD correlation map were very similar to the original results (r = 0.82).

Finally, we replicated our results with a higher resolution parcellation with 100 cortical ROIs (Schaefer et al., 2018). Consistent with our main results, we found that pMFM yielded more realistic simulated FC and FCD in the test set (Figure S6) compared with our previous study (Wang et al., 2019). Across all 10 best parameter sets from the validation set, noise amplitude increased from sensory-motor to association (limbic, control and default) networks, while external input exhibited the opposite direction. In 8 of the 10 best parameter sets, recurrent connect strength increased from sensory-motor to association (limbic, control and default) networks, thus again suggesting potential degeneracy (Figure S7).

In the Schaefer parcellation, time-varying amplitude of sensory-motor time courses tracks switching behavior in time-varying functional connectivity (Figures S8 and S9). Causal perturbation analysis also confirmed that sensory-motor regions appeared to drive transitions in FCD (Figure S9). Both simulated and empirical FCD-STD correlation maps were correlated with PVALB-SST gene expression maps (Table S1). Both correlations were significant under the spin-test and random gene-pair tests. The simulated, but not the empirical, FCD-STD correlation maps were correlated with the first principal component of gene expression.

## Discussion

By incorporating anatomical and functional gradients into the parameterization of local circuit properties, the resulting large-scale circuit model generated realistic time-averaged (static) and time-varying (dynamic) properties of large-scale spontaneous brain activity. Both empirical and simulated fMRI data exhibited multi-stable properties, in which there was spontaneous switching between a high coherent state and a low coherent state. The multi-stability was tracked by time-varying amplitude of regional fMRI signals. By performing causal perturbations of the large-scale circuit model, we demonstrated that spontaneous amplitude fluctuations of sensory-motor regions were a driver of the observed switching behavior. Furthermore, the relationship between regional fMRI amplitude and functional connectivity dynamics was also associated with PVALB-SST and the first principal component of gene expression, suggesting that heterogeneity in excitation-inhibition balance might shape multi-stability in FC dynamics.

### Anatomical and functional gradients contribute to spontaneous brain dynamics

Previous studies have proposed a dominant gradient of cortical organization with sensory-motor and association regions at opposing ends (Huntenburg et al., 2018). Supporting this idea of a dominant axis, many studies have emphasized similarities among gradients estimated from diverse sources, including resting-state FC principal gradient, T1w/T2w myelin estimate, gene expression data, functional task activation and computational modeling (Margulies et al., 2016; Huntenburg et al., 2017; Burt et al., 2018; Wang et al., 2019; Gao et al., 2020). Yet, there are clear differences among the gradients and a growing number of studies have suggested dissociations among multiple spatially similar gradients (Paquola et al., 2019; Shafiei et al., 2020; Valk et al., 2020). Here, we showed that by parameterizing local circuit parameters with both anatomical (T1w/T2w) and functional (FC) gradients, the resulting mean field model was able to generate dramatically more realistic static FC and FC dynamics than either gradient alone (Figure 3).

The optimized mean field model exhibited opposing gradient directions across local circuit parameters (Figure 4). Across all top ten parameter sets, noise amplitude increased from sensory-motor to association cortex, while external input decreased from sensory-motor to association cortex. The higher external input in sensory-motor regions might reflect the flow of sensory information from the external environment via subcortical relays. In the case of the recurrent connection strength, nine of the ten best parameter sets exhibited increasing values from sensory-motor to association cortex, but one parameter set exhibited the opposite direction. Thus, recurrent connection strength might exhibit potential degeneracies in mean field models, thus explaining contradictions in the literature (Demirtas et al., 2019; Wang et al., 2019).

### Multi-stability in spontaneous brain dynamics

The spontaneous ebb and flow observed in FC dynamics is an intriguing property that has fascinated the field (Allen et al., 2014; Hansen et al., 2015; Wang et al., 2016; Liegeois et al., 2017; Vidaurre et al., 2017; Reinen et al., 2018). As shown in Figure 5A, there are periods of brain activity with strong coherent FC and periods with incoherent FC. We found that the coherent FC state was characterized by larger fMRI signal amplitude across brain regions, while the incoherent FC state was characterized by smaller fMRI signal amplitude (Figure 5). Intriguingly, transitions in the regional amplitude of sensory-motor regions appeared to track switching behavior in FC dynamics (Figure 6). Perturbations of the mean field model suggests that this relationship might be causal.

Regional fMRI amplitude has been previously linked with the differential expression of PVALB and SST across the cortex (Anderson et al., 2020a). PVALB and SST interneurons preferentially target perisomatic regions and dendrites of pyramidal cells respectively, and are thought to regulate synaptic outputs and inputs respectively (Wang et al., 2004). Thus the spatially heterogeneous distribution of PVALB and SST interneurons (Kim et al., 2017) might modulate regional neural signal amplitude (Anderson et al., 2020a). Here, we found that PVALB-SST gene expression map correlates with the spatial distribution of sensory-motor drivers whose time-varying amplitude tracks functional connectivity dynamics (Figure 7).

However, we note that this association cannot be solely attributed to PVALB-SST given that the gradients of PAVLB-SST expression are embedded within a broader pattern of gene expression variation across the cortex (Burt et al., 2018; Anderson et al., 2020b). Indeed, the spatial distribution of sensory-motor drivers were also correlated with the first principal component of cortical genes (Figure 7). The first gene principal component has been shown to strongly correlate with the spatial distribution of genes coding for different excitatory and inhibitory neurons (Burt et al., 2018), which might reflect spatial heterogeneity in excitation-inhibition balance (Wang, 2020). Overall, this suggests a potential link between FC dynamics and heterogeneity in excitation/inhibition balance across the cortex.

## Methods

### Data

We considered 1052 participants from the Human Connectome Project (HCP) S1200 release (Van Essen et al., 2013). All participants were scanned on a customized Siemens 3T Skyra using a multi-band sequence. Four resting-state fMRI (rs-fMRI) runs were collected for each participants in two sessions on two different days. Each rs-fMRI run was acquired with a repetition time (TR) of 0.72s at 2mm isotropic resolution and lasted for 14.4 min. The diffusion imaging consisted of 6 runs, each lasting approximately 9 minutes and 50 seconds. Diffusion weighting consisted of 3 shells of b = 1000, 2000, and 3000 s/mm2 with an approximately equal number of weighting directions on each shell. Details of the data collection can be found elsewhere (Van Essen et al., 2013). The 1052 subjects were randomly divided into training (N=351), validation (N=350) and test (N=351) sets.

### Preprocessing

Details of the HCP preprocessing can be found in the HCP S1200 manual. We utilized rs-fMRI data, which had already been projected to fsLR surface space, denoised with ICA-FIX and smoothed by 2mm. For each run of each participant. the fMRI data was averaged within each Desikan-Killiany (Desikan et al., 2006) ROI to generate a 68 x 1200 matrix. Each 68 x 1200 matrix was used to compute 68 x 68 FC matrix by correlating the time courses among all pairs of time courses. The FC matrices were then averaged across runs of participants within the training (or validation or test) set, resulting in a group-averaged training (or validation or test) FC matrix.

Functional connectivity dynamics (FCD) was computed as follows. For each run of each participant, FC was computing within each of 1118 sliding windows. The length of each sliding window was 83 time points (60 seconds) as recommended by previous studies (Leonardi and Van De Ville, 2015; Liegeois et al., 2017). We note that our results were robust to window length (see “Control analysis” in the Results section). Each sliding window FC matrix was then vectorized by only considering the upper triangular entries. The vectorized FCs were correlated with each other generating a 1118 x 1118 FCD matrix.

In the case of diffusion MRI, generalized Q-sampling imaging (GQI) was used to reconstruct the white matter pathways, allowing for complex diffusion fiber configurations and streamline tractography (van den Heuvel and Sporns, 2011). A 68 x 68 structural connectivity (SC) matrix was generated for each subject, where each entry corresponded to the number of streamlines between two ROIs. To generate a group-level SC matrix, a thresholding procedure was employed to remove false positives. More specifically, if less than 50% of participants had a non-zero value in a particular entry in the SC matrix, then the entry is set to zero in all individual-level SC matrices. For each SC entry, the number of streamlines was averaged across participants with non-zero streamlines. Separate group-level SC matrices were computed for the training, validation and test sets.

### Dynamic mean field model (MFM)

The MFM was derived by the mean-field reduction of a detailed spiking neuronal network model (Deco et al., 2013). For each cortical ROI, the neural activity obeys the following nonlinear stochastic differential equations:

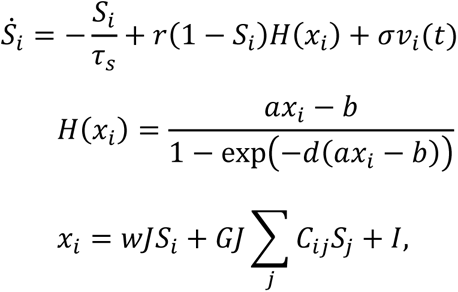

where *S*_*i*_, *H*(*x*_*i*_) and *x*_*i*_ denote the average synaptic gating variable, population firing rate and total input current of the *i*-th cortical ROI. The total input current *x*_*i*_ is the superposition of three inputs. The first input, the intra-regional input, is controlled by the recurrent connection strength *w*. The second input, the inter-regional input, is controlled by the SC matrix (*C*_*ij*_ is the SC between regions *i* and *j*), as well as a global scaling factor *G*. The third input is the external input current *I*, which might include inputs from subcortical relays. Following previous studies (Deco et al., 2013; Wang et al., 2019), the synaptic coupling *J* was set to 0.2609 (*nA*). The parameter values of the input-output function *H*(*x*_*i*_) were set to *a* = 270(*n*/*C*), *b* = 108(*Hz*) and *d* = 0.154(*s*). The kinetic parameters for synaptic activity were set to *r* = 0.641 and *τ*_*s*_ = 0.1(*s*). *v*_*i*_(*t*) is uncorrelated standard Gaussian noise and the noise amplitude is controlled by *σ*.

The simulated neural activities *S*_*i*_ are fed to the Balloon-Windkessel hemodynamic model (Stephan et al., 2007; Heinzle et al., 2016) to simulate the fMRI BOLD signals for each ROI. The equations and parameters are exactly the same as our previous study (Wang et al., 2019). More specifically, the MFM and hemodynamic model were simulated using Euler’s integration with time step of 10ms. The starting values of *S*_*i*_ in the MFM were randomly initialized. Simulation length for the fMRI signals was 16.4 min. The first 2 minutes of the fMRI signals were discarded and the time series were downsampled to 0.72s to have the same temporal resolution as the empirical fMRI signals in the HCP. The simulated fMRI signals could then be used to generate simulated FC and FCD matrices.

### Parametric Mean Field Model (pMFM)

In our previous study (Wang et al., 2019), the recurrent connection strength *w*, external input current *I*, global constant *G* and noise amplitude *σ* were optimized by fitting to static FC. The recurrent connection strength *w* and external input current *I* were allowed to vary independently across cortical ROIs, while *G* and *σ* were assumed to be constant. On the other hand, (Demirtas et al., 2019) parameterized the recurrent connection strengths with the T1w/T2w myelin map.

In this study, recurrent connection strength *w*, external input current *I* and noise amplitude *σ* were allowed to vary across brain regions, while *G* was kept as a constant. Instead of allowing *w*, *I* and *σ* to vary independently (Wang et al., 2019), we parameterized *w*, *I* and *σ* as linear combinations of group-level T1w/T2w myelin maps (Glasser and Van Essen, 2011) and the first principal gradient of functional connectivity (Margulies et al., 2016):

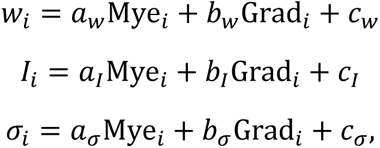

where *w*_*i*_, *I*_*i*_ and *σ*_*i*_ denoted the recurrent connection strength, external input current and noise amplitude respectively of the *i*-th cortical region. Mye_*i*_ and Grad_*i*_ were the average values of the T1w/T2w myelin map and the first FC principal gradient within the *i*-th cortical ROI. Both T1w/T2w myelin maps and first principal gradient of functional connectivity were computed from the HCP training set. Therefore, there are a total of 10 unknown parameters: G and linear coefficients (*a*_*w*_, *b*_*w*_, *c*_*w*_, *a*_*I*_, *b*_*I*_, *c*_*I*_, *a*_*σ*_, *b*_*σ*_, *c*_*σ*_). These unknown parameters were be estimated from the HCP training set (see next section).

### Cost function to minimize disagreement with empirical static FC and FCD

The 10 unknown parameters in the pMFM were estimated by maximizing fit to static FC and FCD in the HCP training set. For a particular set of parameters, the pMFM could be used to generate simulated FC and FCD matrices. The agreement between the simulated and empirical static FC matrices was defined as the Pearson’s correlation (r) between the z-transformed upper triangular entries of the two matrices. Larger r indicates more similar static FC.

The disagreement between the simulated and empirical FCD matrices was defined as the Kolmogorov–Smirnov (KS) distance between the probability distribution functions (pdfs) constructed from the upper triangular entries of the two matrices (Hansen et al., 2015). A smaller KS distance indicated more similar FCD. To optimize fit to both static FC and FCD, an overall cost was defined as (1 - r) + KS. Thus lower cost implies better fit to static FC and FCD.

To minimize the cost function in the training set, we seek to compute an “average” FCD matrix. We note that FCD matrices could not be directly averaged across rs-fMRI runs and participants because there was no temporal correspondence across runs during the resting-state. Because the goal here was to compute the KS distance, we simply averaged the pdfs from the FCD matrices all the runs of all participants within the training set, which we referred to as average FCD pdf. When evaluating KS distance in the validation and test sets, average FCD pdfs were also computed using the same approach.

### Optimization procedure

To optimize the cost function, we considered three algorithms: covariance matrix adaptation evolution strategy (CMA-ES; Hansen, 2006), self-organising migrating algorithm (SOMA; Davendra and Zelinka, 2016) and hyperparameter optimization using radial basis functions and dynamic coordinate search (HORD; Ilievski et al., 2017).

Given a particular random initialization of the 10 unknown parameters, the three algorithms (CMA-ES, SOMA, HORD) were applied to the HCP training set. Each algorithm was iterated 500 times, generating 500 candidate parameter sets. This procedure was repeated 10 times, yielding 5000 candidate parameter sets. For each algorithm, the 5000 candidate parameter sets were evaluated in the validation set to obtain top 10 candidate parameter sets. Across the three algorithms, CMA-ES performed the best in the validation set (Figure S1), so this study focused on CMA-ES.

The top 10 candidate parameter sets from CMA-ES were then applied to the HCP test set SC. For each parameter set, 1000 simulations were performed, yielding 1000 simulated static FC and FCD matrices. The 1000 simulated FC and FCD pdfs were then averaged, yielding an average simulated FC and an average simulated FCD pdf. Pearson’s correlation was then computed between the average simulated FC and the average empirical FC from the HCP test set. Similarly, KS statistics was computed between the average simulated FCD pdf and the average empirical FCD pdf from the HCP test set.

### Statistical test of correlation between first derivatives of FCD mean and SW-STD

To quantify the correspondence between FCD mean and SW-STD (Figure 5), correlation between the first derivative of the FCD mean time course and the first derivative of the SW-STD time course was computed for each cortical region (Figure 6). To compute the statistical significance of the correlations, fMRI runs were permuted across participants. For each ROI, the FCD-STD correlations were recomputed and averaged across runs and participants, yielding a single null correlation value. This permutation procedure was repeated 10000 times, so that a null distribution of correlations was obtained for each ROI.

### Causal perturbations of pMFM

To more directly link sensory-motor regions with FCD, we tested whether perturbation of sensory-motor regions can “kick” the system from an incoherent FCD state to a coherent FCD state. Among 1000 random simulations of the pMFM, time segments in the incoherent (low FCD mean) state lasting for at least 200 contiguous fMRI timepoints (TRs) were selected, yielding 300 time segments. Low FCD mean was defined as being less than 0.6.

Perturbation was applied to the neural signals (synaptic gating variable *S*_*i*_) of the top 5 regions whose SW-STD correlated with FCD (Figure 6B). We note that during the incoherent state, the values of the synaptic gating variables could be low or high. To increase the amplitude of the neural signals, we would decrease (or increase) the synaptic gating variables if they were high (or low). More specifically, let *S*_*max*_ and *S*_*min*_ be the maximum and minimum synaptic gating variable values across all cortical regions. When neural signal was low, we set *S*_*t*+*δt*_ = *S*_*t*_ + 0.8 (*S*_*max*_ − *S*_*t*_), where *δt* corresponded to the resolution of the simulations, which is 0.01 seconds in the current study. When neural signal was high, we set *S*_*t*+*δt*_ = *S*_*t*_ − 0.8 (*S*_*t*_ − *S*_*min*_). The perturbations was applied for 72 iterations, corresponding to 1 TR in the simulated fMRI signal.

### Gene expression analysis

Publicly available human gene expression data from six postmortem donors (1 female), aged 24–57 years (42.5 ± 13.4) were obtained from the Allen Institute (Hawrylycz et al., 2012). Processing followed the pipeline from Anderson and colleagues (Anderson et al., 2020a; https://github.com/HolmesLab/2020_NatComm_interneurons_cortical_function_schizophrenia), yielding 17,448 brain-expressed genes and 1683 analyzable cortical samples. Our analyses in turn focused on 2413 brain-specific genes (Genovese et al., 2016; Burt et al., 2018). Z-normalized gene expression values of parvalbumin (PVALB) and somatostatin (SST) were averaged within each cortical region and the difference was computed. The FCD-STD correlation maps (Figure 6) were correlated with the PVALB-SST spatial map (Figure 7).

To establish statistical significance, we considered two approaches. First, we considered the spin test. The parcellations were randomly rotated. For each rotated parcellation, we re-computed the PVALB-SST difference and correlated the resulting gene expression maps with the FCD-STD correlation maps, yielding a single null correlation value. This was repeated 1000 times yielding a complete null distribution.

To test the specificity of PVALB-SST, we performed a random-gene-pair tests. A random pair of genes was selected from the 2413 brain-specific genes (Burt et al., 2018). Gene expression difference between the random gene pairs was computed and correlated with the STD-FCD correlation maps generating a null correlation value. This was repeated 10,000 times yielding a complete null distribution.

### Code and data availability

This study followed the institutional review board guidelines of corresponding institutions. The HCP diffusion MRI, rs-fMRI and T1w/T2w data are publicly available (https://www.humanconnectome.org/). The code used in this paper is publicly available at https://github.com/ThomasYeoLab/CBIG/tree/master/stable_projects/fMRI_dynamics/Kong2021_pMFM. The code was reviewed by one of the co-authors (SZ) before merging into the GitHub repository to reduce the chance of coding errors.

## Acknowledgements

This work was supported by the Singapore National Research Foundation (NRF) Fellowship (Class of 2017) and the National University of Singapore Yong Loo Lin School of Medicine (NUHSRO/2020/124/TMR/LOA). Any opinions, findings and conclusions or recommendations expressed in this material are those of the authors and do not reflect the views of the National Research Foundation, Singapore. Our computational work was partially performed on resources of the National Supercomputing Centre, Singapore (https://www.nscc.sg). Our research also utilized resources provided by the Center for Functional Neuroimaging Technologies, P41EB015896 and instruments supported by 1S10RR023401, 1S10RR019307, and 1S10RR023043 from the Athinoula A. Martinos Center for Biomedical Imaging at the Massachusetts General Hospital. Data were in part provided by the Human Connectome Project, WU-Minn Consortium (Principal Investigators: David Van Essen and Kamil Ugurbil; 1U54MH091657) funded by the 16 NIH Institutes and Centers that support the NIH Blueprint for Neuroscience Research; and by the McDonnell Center for Systems Neuroscience at Washington University.

## Supplementary Results

**Figure S1.**
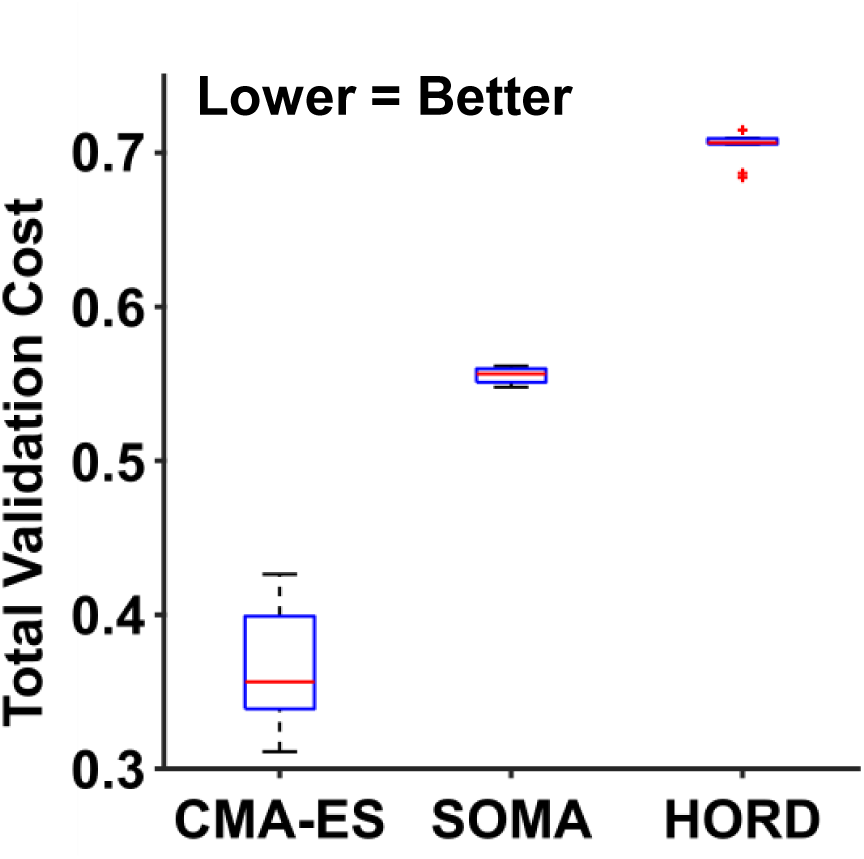
Comparison of three different algorithms: covariance matrix adaptation evolution strategy (CMA-ES; Hansen, 2006), self-organising migrating algorithm (SOMA; Davendra and Zelinka, 2016) and hyperparameter optimization using radial basis functions and dynamic coordinate search (HORD; Ilievski et al., 2017) in the HCP validation set. Each algorithm was run on the training set generating 5000 candidate sets of model parameters. The 5000 candidate sets were evaluated in the validation set. The top 10 candidate sets from each algorithm (based on the validation set) are shown in this plot. Thus, CMA-ES performs the best among the three algorithms in the validation set. Box plots utilized default Matlab parameters, i.e., box shows median and inter-quartile range (IQR). Whiskers indicate 1.5 IQR. Red crosses represent outliers.

**Figure S2.**
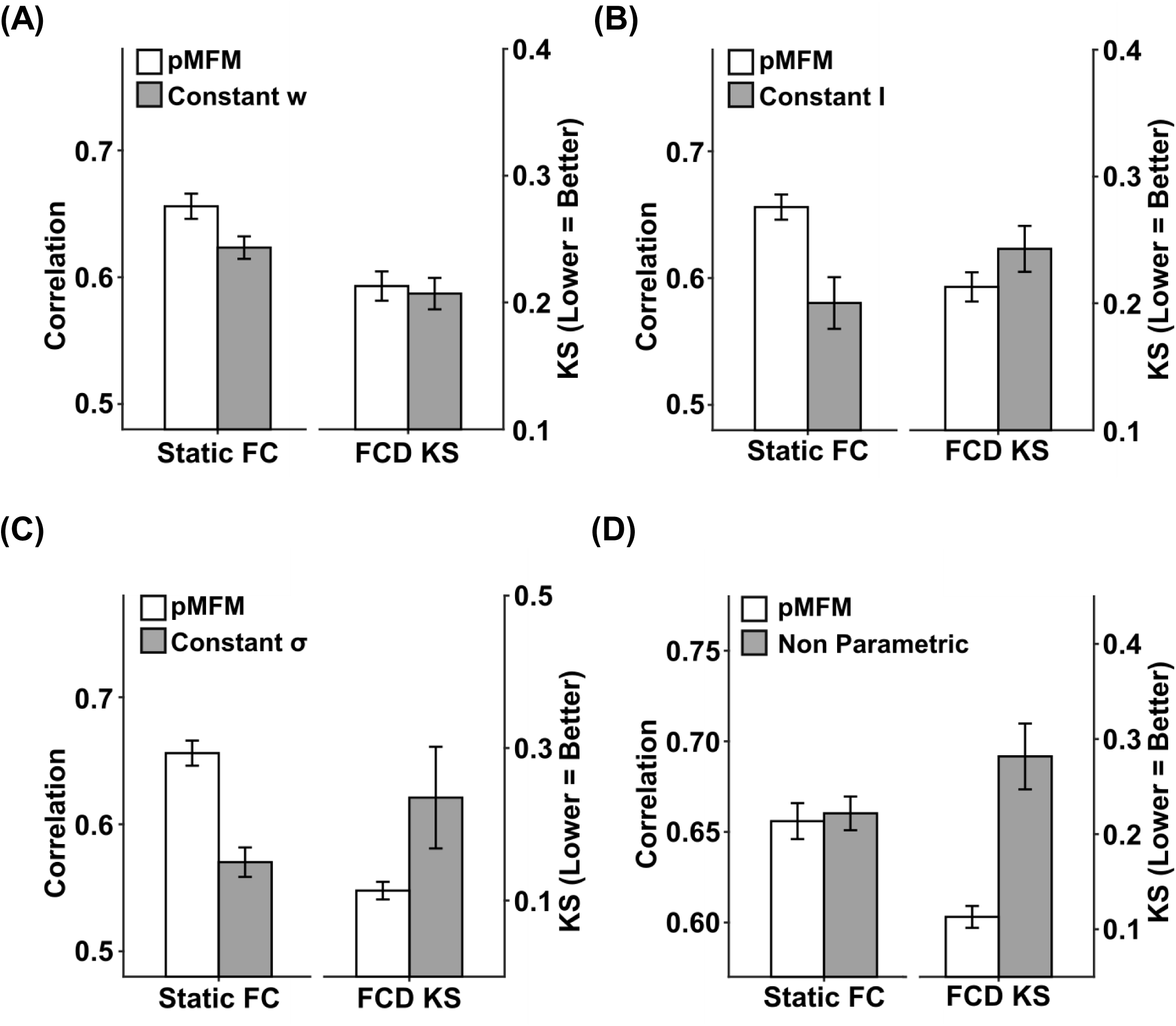
Comparison between the original pMFM (main text) and (A) constraining recurrent connection strength *w* to be constant across ROIs, (B) constraining external input *I* to be constant across ROIs, (C) constraining noise amplitude *σ* to be the same across ROIs, and (D) allowing local circuit parameters to vary independent (i.e., not parameterized by anatomical and/or functional gradients). Across all panels, agreement between simulated and empirical static FC was measured using Pearson’s correlation, while disagreement between simulated and empirical FCD was measured using KS distance. Across all analyses, top ten model parameter sets were selected from the validation set and applied to the test set. The error bars correspond to standard error across the 10 parameter sets. Across all four panels, the original pMFM yielded the best results.

**Figure S3.**
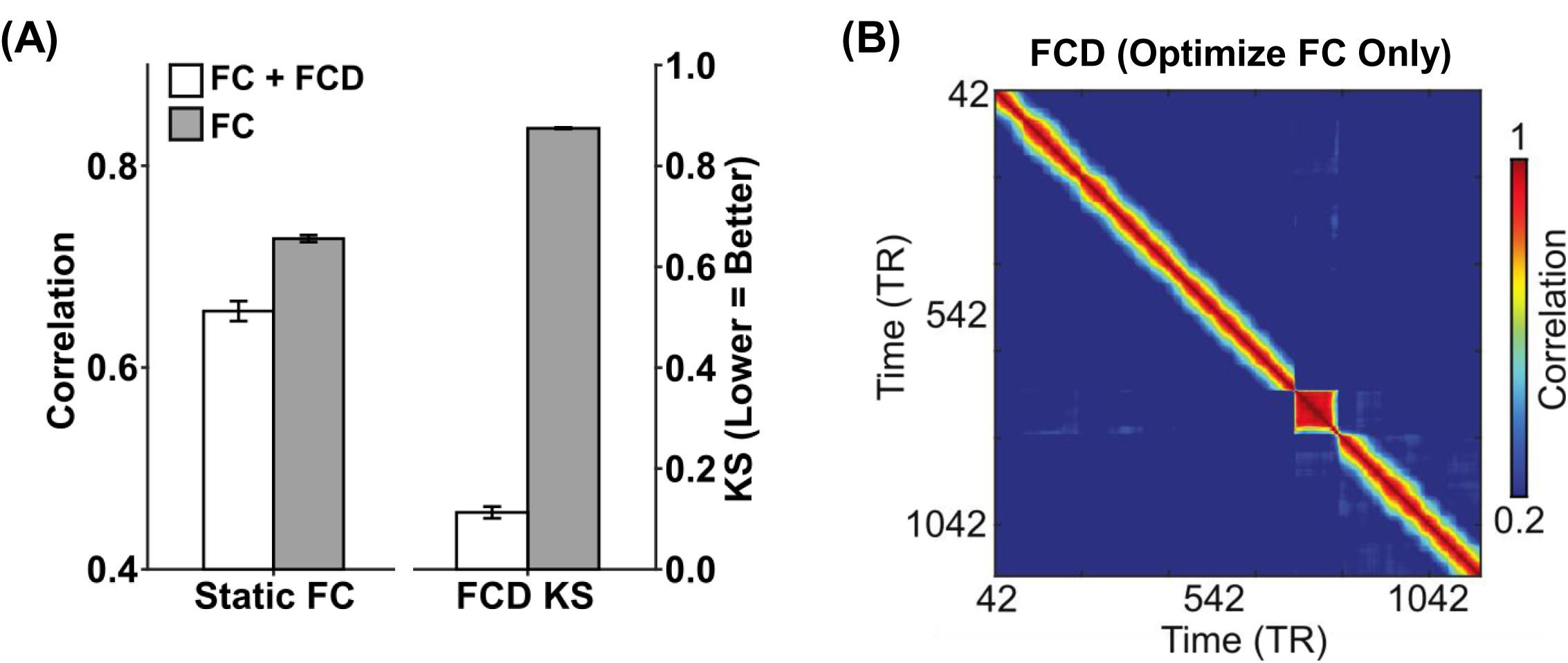
Comparison between the original pMFM (optimized using both static FC and FCD) and pMFM optimized using only static FC. (A) Agreement (Pearson’s correlation r) between simulated and empirically observed static FC, as well as disagreement (KS distance) between simulated and empirically observed FCD. (B) Simulated FCD from the pMFM optimized only using static FC. The simulated FCD was a lot less realistic than the original pMFM (Figure 2B). In terms of KS distance, there is a large improvement when optimizing both static FC and FCD (KS = 0.12 versus 0.88). However, when optimizing only static FC, the resulting simulated static FC was only slightly better than the original pMFM (r = 0.74 versus 0.66). This suggests that the goals of generating realistic static FC and FCD were not necessarily contradictory. We note that across all analyses, top ten model parameter sets were selected from the validation set and applied to the test set. The error bars correspond to standard error across the 10 parameter sets.

**Figure S4.**
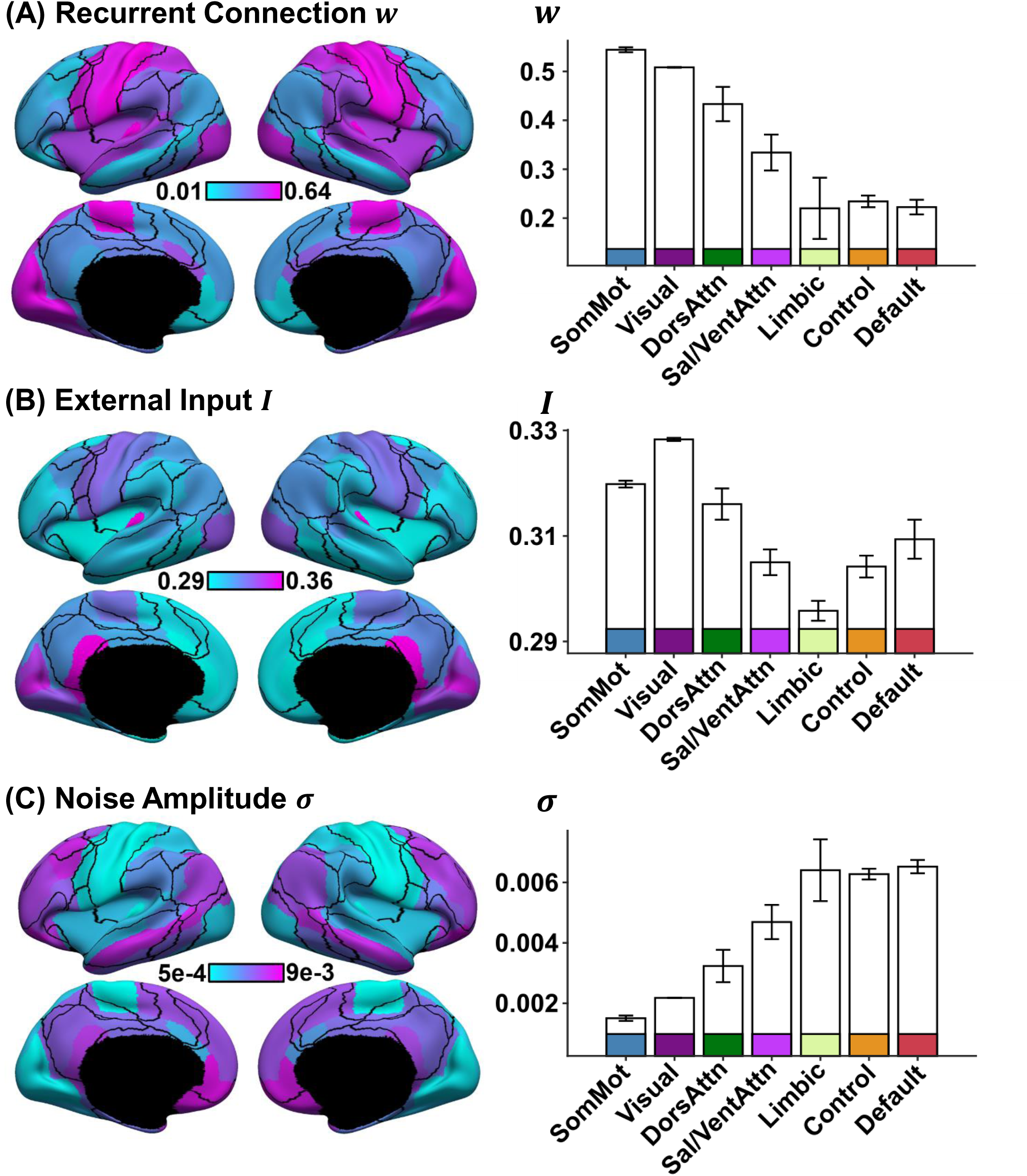
For one of the top ten parameter sets, recurrent connection strength exhibited the opposite direction from the remaining nine parameter sets. The layout of this figure is the same as Figure 4. (A) Strength of recurrent connection *w* in 68 Desikan-Killiany cortical ROIs (left) and seven resting-state networks (right). (B) Strength of external input *I* in 68 Desikan-Killiany cortical ROIs (left) and seven resting-state networks (right). (C) Strength of noise amplitude *σ* in 68 Desikan-Killiany cortical ROIs (left) and seven resting-state networks (right). The bars represent the mean values across regions within each network. The error bars show the standard error across regions within each network. Noise amplitude increased from sensory-motor to association (limbic, control and default) networks. On the other hand, external input current and recurrent connection strength decreased from sensory-motor to association networks.

**Figure S5.**
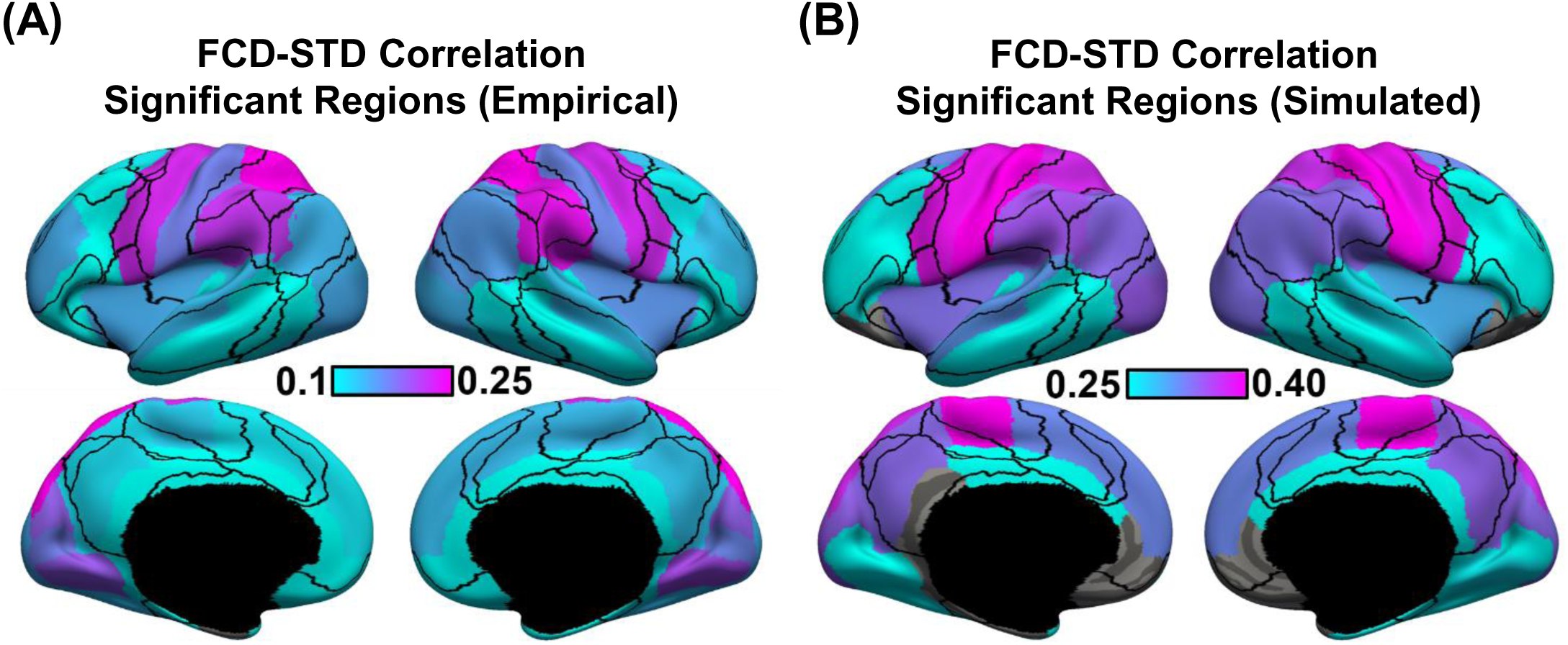
Sensory-motor regions drive sharp transitions in functional connectivity dynamics (FCD). (A) FCD-STD correlations obtained by correlating the first derivative of the FCD mean time course and the first derivative of the SW-STD time course of each cortical region. These correlations were performed for each HCP test participant and averaged across all runs and participants. Regions that survived a false positive rate of q<0.05 are shown in the brain map. (B) Same as panel A but simulated from pMFM using the best model parameters from the validation set and structural connectivity from the test set. Regions that survived a false positive rate of q<0.05 are shown in the brain map.

**Figure S6.**
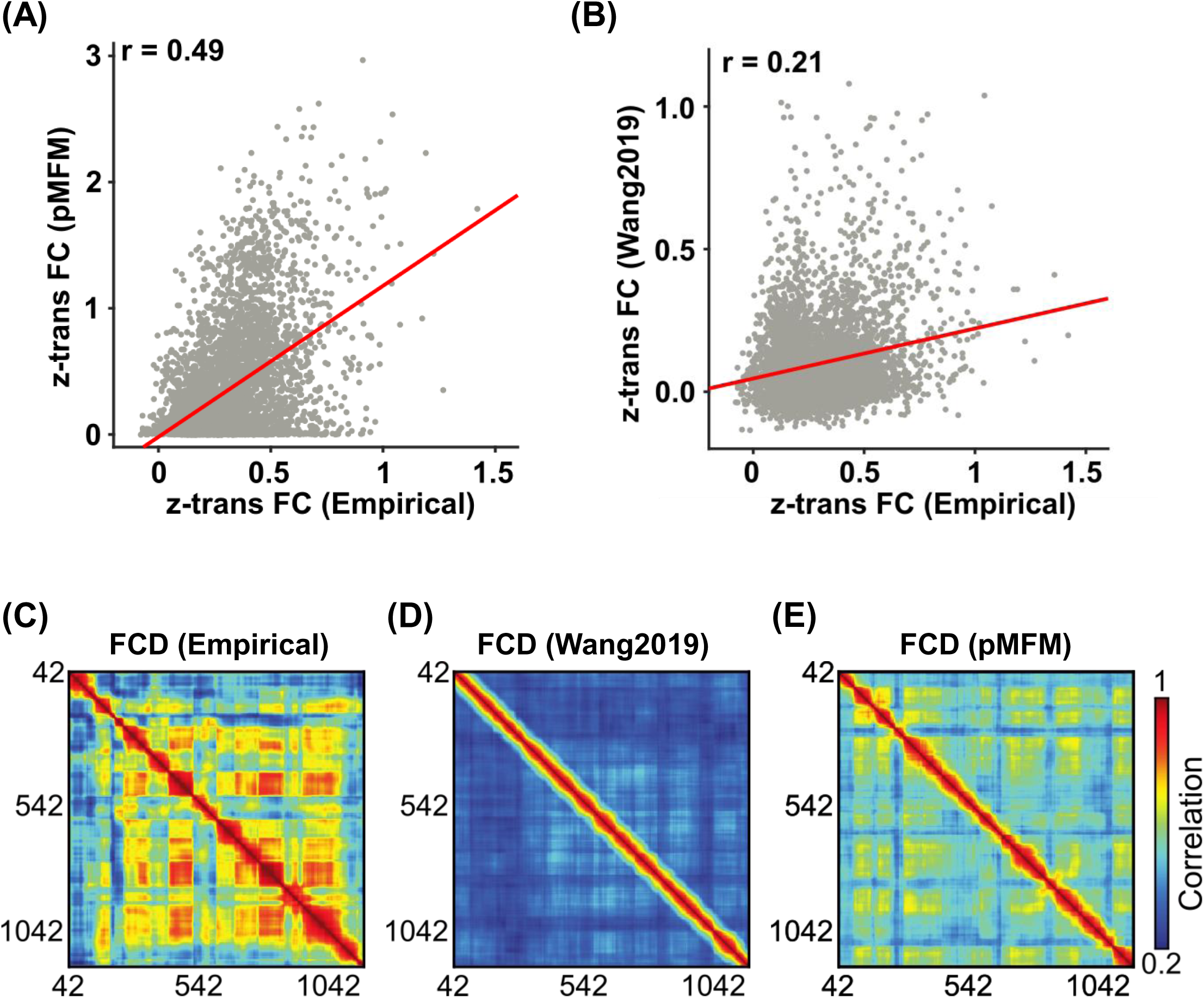
Parametric mean field model (pMFM) generates more realistic static functional connectivity (FC) and functional connectivity dynamics (FCD) than the previous spatially heterogeneous MFM (Wang et al., 2019) in the 100-region Schaefer parcellation. This figure is similar to Figure 2 but utilizes the 100-region Schaefer parcellation. (A) Agreement (Pearson’s correlation) between empirically observed and pMFM-simulated static FC. (B) Agreement (Pearson’s correlation) between empirically observed and simulated static FC from Wang 2019. (C) Empirical FCD from a participant from the HCP test set. (D) Simulated FCD from the pMFM using the best model parameters from the validation set using structural connectivity (SC) from the test set. (E) Simulated FCD generated by the previous spatially heterogeneous MFM (Wang et al., 2019).

**Figure S7.**
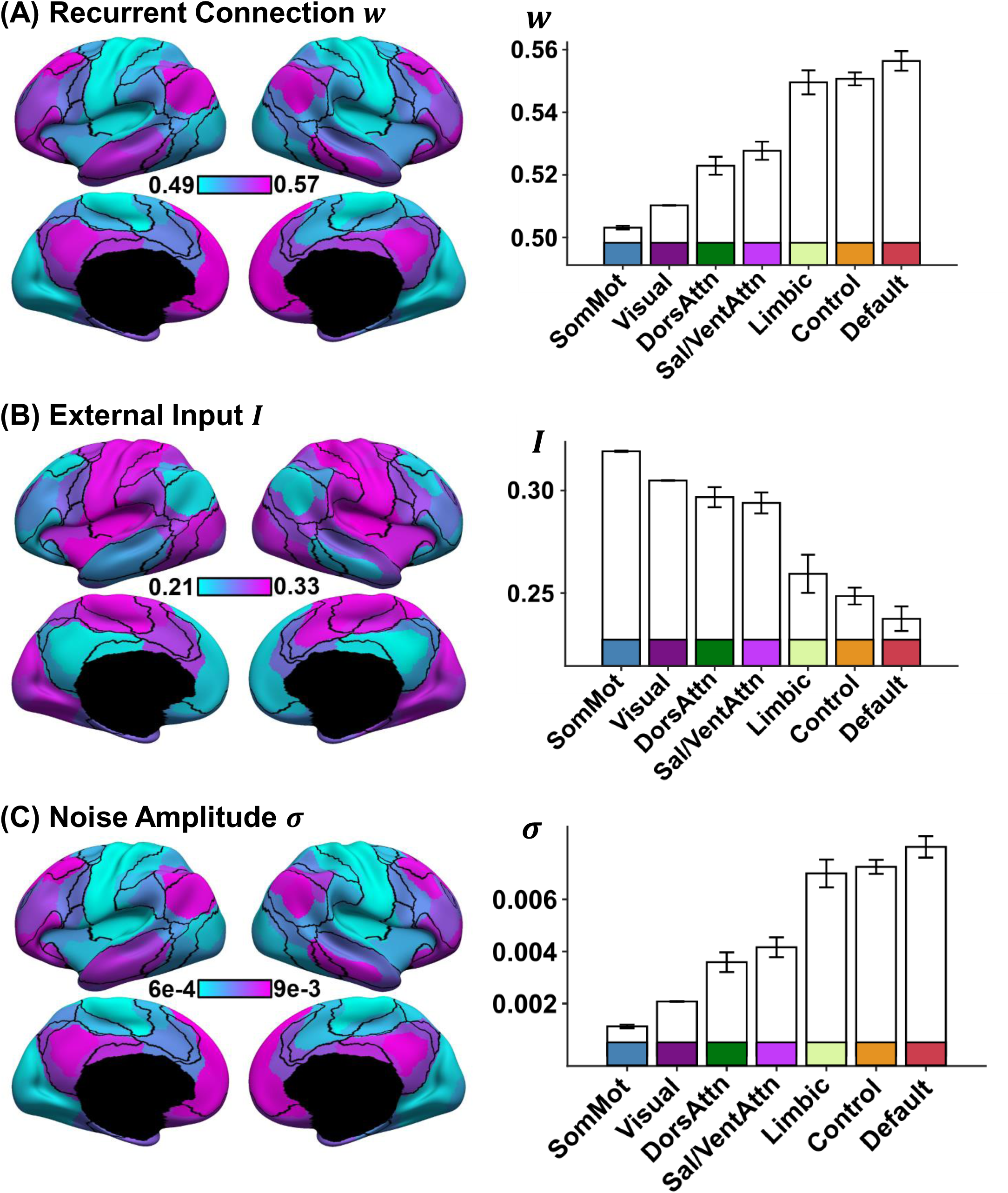
Spatial distribution of recurrent connection strength *w*, external input current *I*, and noise amplitude *σ*, and their relationships with resting-state networks in the 100-region Schaefer parcellation. This figure is similar to Figure 4 but utilizes the 100-region Schaefer parcellation. (A) Strength of recurrent connection *w* in 100 Schaefer cortical ROIs (left) and seven resting-state networks (right). (B) Strength of external input *I* in 100 Schaefer cortical ROIs (left) and seven resting-state networks (right). (C) Strength of noise amplitude *σ* in 100 Schaefer cortical ROIs (left) and seven resting-state networks (right). The bars represent the mean values across regions within each network. The error bars show the standard error across regions within each network. Recurrent connection strength and noise amplitude increased from sensory-motor to association (limbic, control and default) networks. On the other hand, external input current was the highest in sensory-motor networks and decreased towards the default network.

**Figure S8.**
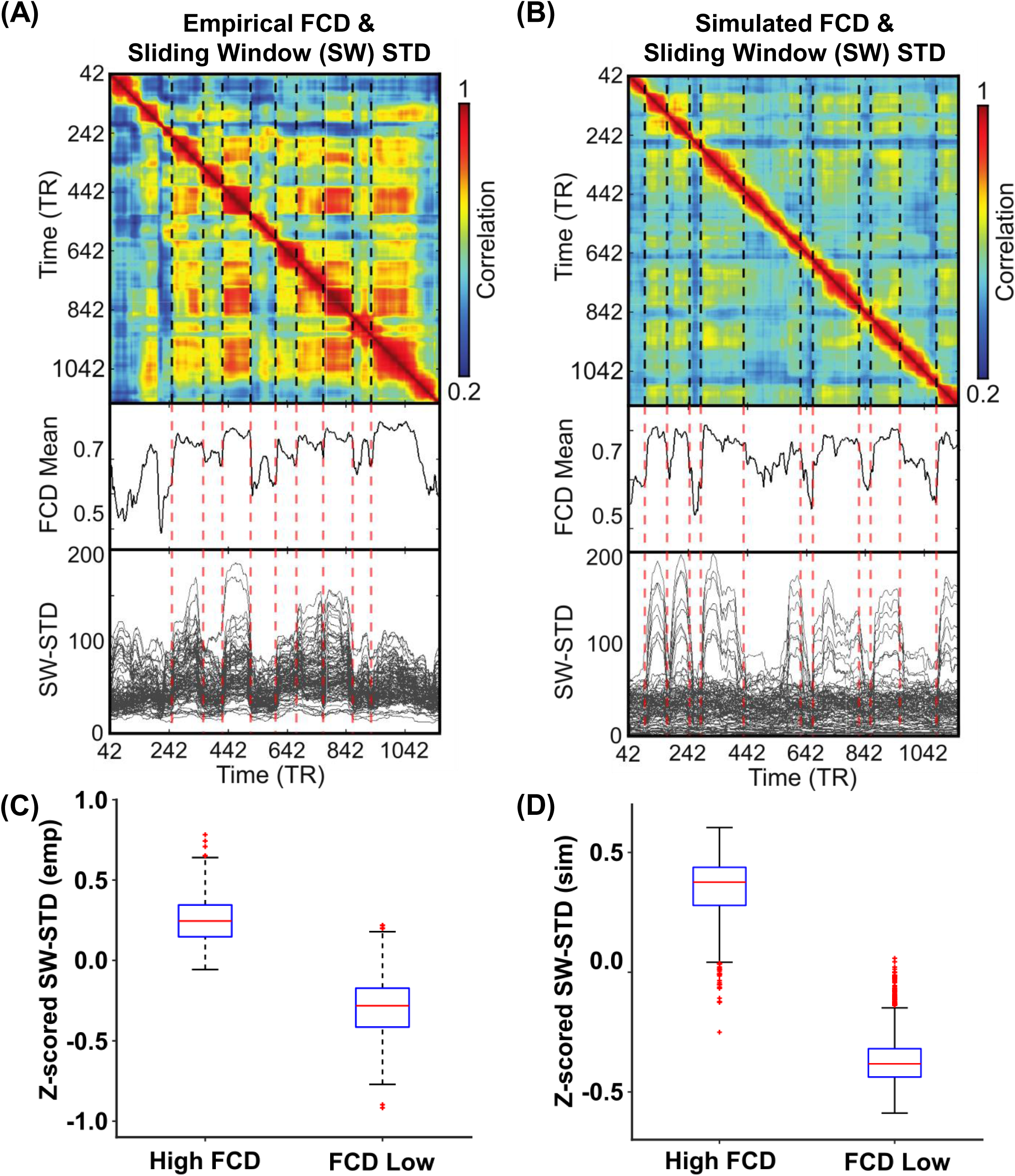
Correspondence between functional connectivity dynamics (FCD) and time-varying amplitude of regional fMRI time courses using the 100-region Schaefer parcellation. This figure is similar to Figure 5 but utilizes the 100-region Schaefer parcellation. (A) Top panel shows empirical FCD matrix of a HCP test participant. The middle panel shows the FCD mean time course obtained by averaging the rows of the FCD matrix from the top panel. The bottom panel shows the standard deviation of each regional fMRI time course within each sliding window (SW-STD). The color of the lines corresponds to the correlation between the first derivative of the FCD mean time course and the first derivative of the SW-STD time courses. Sharp transitions in SW-STD corresponded to sharp FCD transitions (red dashed lines). (B) Same as panel A, but simulated from pMFM using the best model parameters from the validation set and structural connectivity from the test set. (C) SW-STD during coherent (high FCD mean) and incoherent (low FCD mean) states. Boxplots illustrate variation across HCP test participants. Coherent states were characterized by large amplitude (STD) in fMRI signals (p = 4.4e-115). (D) Same as panel C but simulated from pMFM.

**Figure S9.**
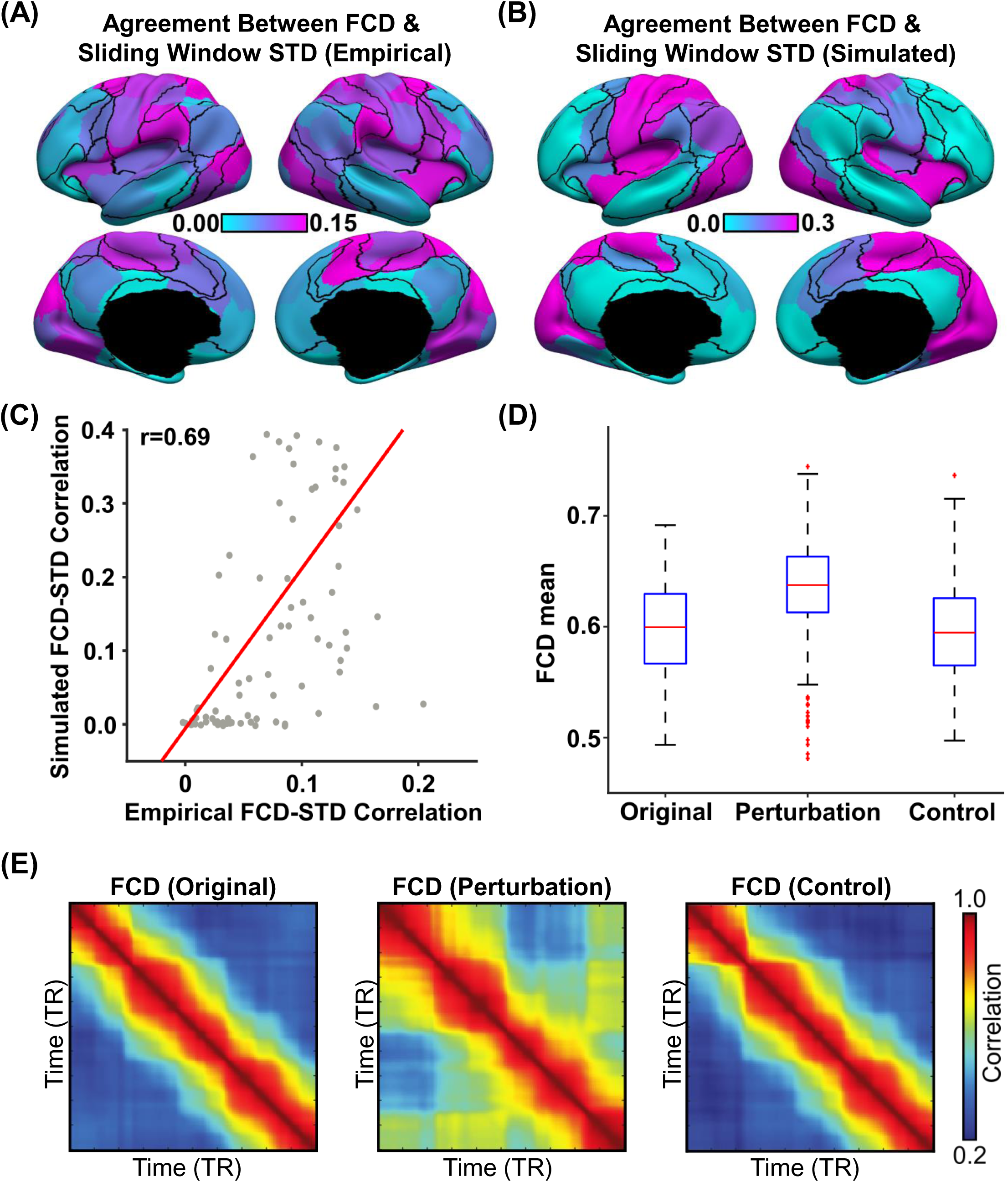
Sensory-motor regions drive sharp transitions in functional connectivity dynamics (FCD) in the 100-region Schaefer parcellation. This figure is similar to Figure 6 but utilizes the 100-region Schaefer parcellation (A) FCD-STD correlations obtained by correlating the first derivative of the FCD mean time course and the first derivative of the SW-STD time course of each cortical region. (B) Same as panel A but simulated from pMFM (C) Correlation between empirical and simulated FCD-STD correlation spatial maps from panels B and C, showing strong correspondence between empirical and simulated results. (D) Casual perturbation of top 5 FCD-STD correlated regions (panel B) during the incoherent state (low FCD mean) led to transition into the coherent state (high FCD mean). As a control analysis, perturbation of the bottom 5 FCD-STD correlated regions (panel B) during the incoherent state (low FCD mean) did not lead to a state change (FCD mean remains low). (E) Example FCD from the perturbation experiments. (Left) original incoherent state. (Middle) perturbation of top 5 FCD-STD correlated regions (sensory-motor drivers). (Right) perturbation of bottom 5 FCD-STD correlated regions.

**Table S1.**
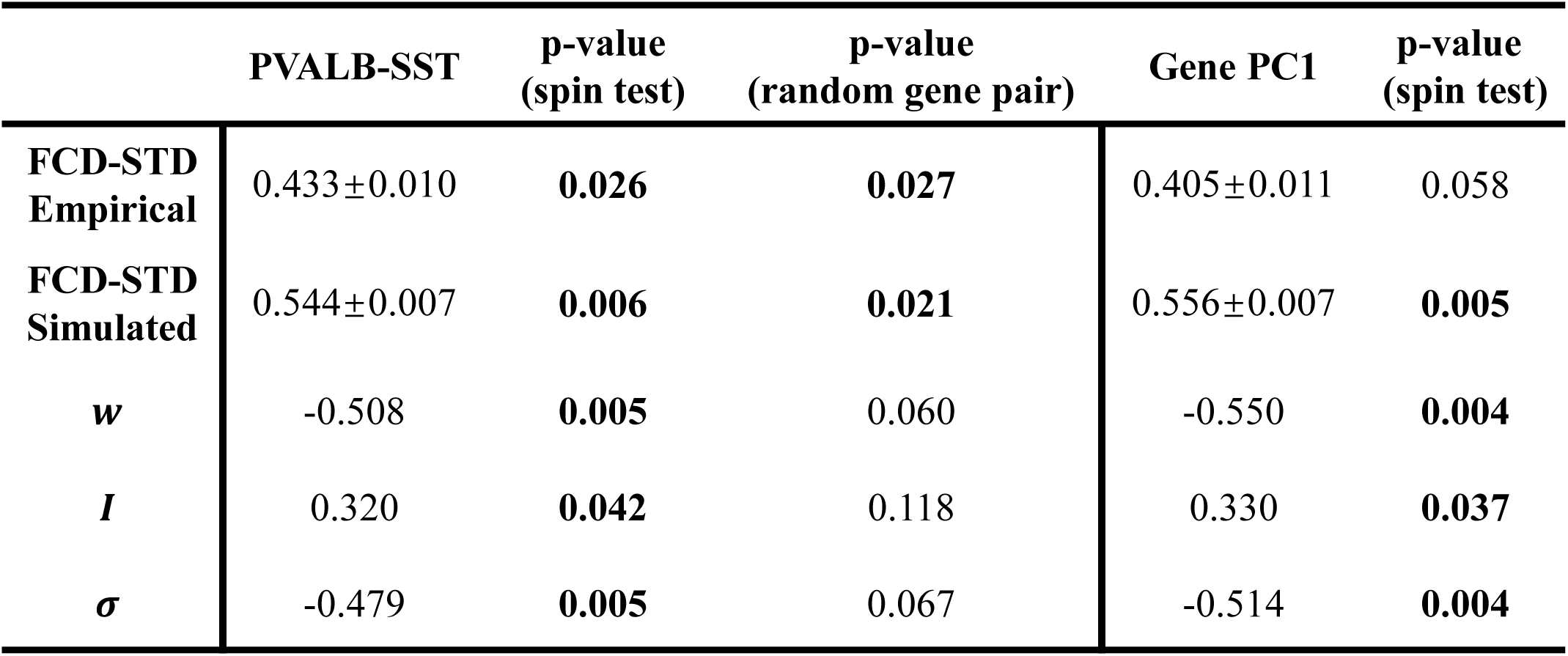
Table of correlations between FCD-STD correlational spatial maps and two gene expression maps: PVALB-SST and first principal component of gene expression (Burt et al., 2018; Anderson et al., 2020b). P values that survived the false discovery rate (q < 0.05) are bolded. Standard deviations reported in the table were obtained by bootstrapping.

|  | PVALB-SST | p-value<br>(spin test) | p-value<br>(random gene pair) | Gene PC1 | p-value<br>(spin test) |
| --- | --- | --- | --- | --- | --- |
| FCD-STD Empirical | 0.433±0.010 | **0.026** | **0.027** | 0.405±0.011 | 0.058 |
| FCD-STD Simulated | 0.544±0.007 | **0.006** | **0.021** | 0.556±0.007 | **0.005** |
| *w* | -0.508 | **0.005** | 0.060 | -0.550 | **0.004** |
| *I* | 0.320 | **0.042** | 0.118 | 0.330 | **0.037** |
| *σ* | -0.479 | **0.005** | 0.067 | -0.514 | **0.004** |

